# Dysfunctional effector programming in Foxp3-ΔE2 Tregs promotes enhanced CD8^+^ T cell responses

**DOI:** 10.64898/2026.04.20.719728

**Authors:** Kristin N. Weinstein, Zoe H. Bishop, Elya A. Shamskhou, Finnegan N. Barry, Gabriela Verdezoto, Marissa de Leon, Joseph R. Albe, Andrew Koval, Baohua Zhou, Phillip P. Domeier, Michael Y. Gerner, Daniel J. Campbell, Steven F. Ziegler

## Abstract

Regulatory T cells (Tregs) prevent autoimmunity through suppressive functions largely programmed by the transcription factor FOXP3. Humans express two major alternatively spliced isoforms of FOXP3: a full-length version containing all coding exons (FOXP3-FL) and a version lacking Exon 2 (FOXP3-ΔE2). Sole FOXP3-ΔE2 expression causes lethal IPEX syndrome, indicating that FOXP3 Exon 2 is essential for full Treg function. Here, we used Foxp3-ΔE2 mice to determine how Foxp3 Exon 2 impacts the functional differentiation of Tregs at homeostasis and during inflammation. Analysis of central and effector Foxp3-ΔE2 Treg populations revealed dysfunctional effector Treg programming, including reduced expression of IL-2 receptor subunits, CTLA-4, and tissue-homing chemokine receptors. During influenza infection, reduced IL-2 regulation by Foxp3-ΔE2 Tregs and decreased Foxp3-ΔE2 Treg localization to the T cell zone during priming contributed to enhanced acute and memory CD8^+^ T cell responses. Thus, dysfunctional effector Treg programming by the Foxp3-ΔE2 isoform impairs suppressive function, resulting in failure to restrain CD8^+^ T cells and aberrant immune responses.

**One Sentence Summary:** Foxp3-ΔE2 expressing regulatory T cells have dysfunctional effector programming which impairs their IL-2 regulation function and co-localization with conventional T cells during priming, thereby enhancing CD8^+^ T cell responses.

**Graphical abstract:** 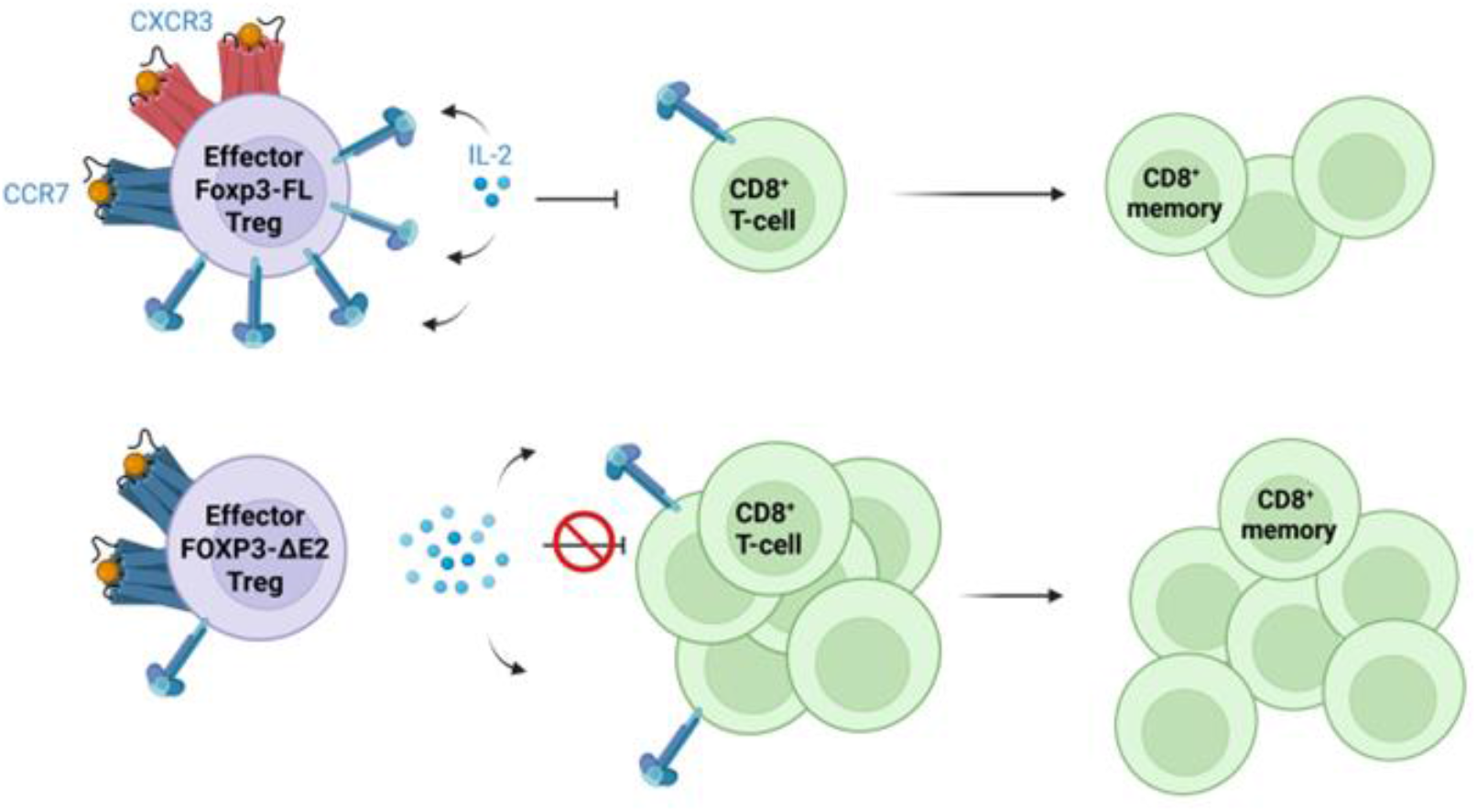

Dysfunctional Foxp3-ΔE2 effector Treg programming includes reduced IL-2Rα and CXCR3 expression and increased CCR7 expression, driving reduced Foxp3-ΔE2 Treg IL-2 sink activity and localization to the T cell zone of the lymph node during influenza viral infection. Together, these defective mechanisms of suppression by Foxp3-ΔE2 Tregs promote CD8^+^ T cell responses during acute infection and memory formation in Foxp3-ΔE2 mice.

## INTRODUCTION

Regulatory T cells (Tregs) are essential for the maintenance of immune homeostasis and prevention of autoimmunity (*1, 2*). Tregs function through a variety of mechanisms, including direct cell-cell contact mediated inhibition by CTLA-4 (*3*), secretion of inhibitory cytokines (*4, 5*), and metabolic disruption of target cells (*6, 7*). Constitutively high cell surface expression levels of the high-affinity IL-2Rα (CD25) allows IL-2 sequestration by Tregs, limiting conventional T cells access to survival and proliferation signaling (*7*). These critical suppressive functions of Tregs are largely programmed by the transcription factor Forkhead Box P3 (FOXP3) (*8–10*).

Tregs in healthy humans have approximately equivalent expression levels of two alternatively spliced isoforms of FOXP3 (*11*). The first isoform contains all 11 coding exons and is referred to as FOXP3-Full Length (FOXP3-FL) while the shorter isoform lacks Exon 2 (FOXP3-ΔE2). FOXP3 Exon 2 encodes binding sites for RORα, RORγt, and hnRNPF, suggesting functional differences between the two isoforms (*12–14*). Although these functional differences remain incompletely defined, sole expression of the FOXP3-ΔE2 isoform drives lethal immune dysregulation, polyendocrinopathy, enteropathy, X-linked (IPEX) syndrome (*15*). Additionally, the FOXP3-ΔE2 isoform is elevated in several autoimmune diseases, including Hashimoto’s thyroiditis (*16*), antineutrophil cytoplasmic antibody-associated vasculitis (*17*), and celiac disease (*18*). These results strongly suggest defects in immune regulation by FOXP3-ΔE2 Tregs.

While rodents only naturally express the Foxp3-FL isoform, the development of a Foxp3-ΔE2 mouse via deletion of the second coding exon of the *Foxp3* gene allows study of Foxp3-ΔE2 expressing Tregs *in vivo* (*15, 19*). In these mice, Foxp3-ΔE2 Tregs skew toward a CD62L^hi^ CD44^lo^ central Treg differentiation state, are outcompeted by Foxp3-FL Tregs in heterozygous Foxp3-FL/ΔE2 mice, and have reduced expression of important functional molecules, including CD25 and CTLA-4. Foxp3-ΔE2 mice also have spontaneously increased conventional T cell activation although they do not develop lethal autoimmunity as observed in humans with sole FOXP3-ΔE2 expression. Interestingly, both murine and engineered human Foxp3-ΔE2 Tregs suppress CD4^+^ Foxp3^-^T cell proliferation *in vitro* (*15, 20, 21*). These findings raised two key questions: (1) whether Foxp3 Exon 2 differentially regulates the biology of central and/or effector Treg populations and (2) which mechanisms of conventional T cell regulation are defective in Foxp3-ΔE2 Tregs.

In this study, we demonstrate that the functional impact of Foxp3 Exon 2 is highly context dependent, profoundly shaping the transcriptional and phenotypic programming of effector Tregs with only a modest impact on central Tregs. These differences included reduced CD25 and CTLA-4 expression and altered tissue-homing chemokine receptor expression.

Dysfunctional effector Treg programming in Foxp3-ΔE2 Tregs results in defective regulation of CD8^+^ T cell responses during influenza viral infection, leading to enhanced memory formation following infection. Impaired IL-2 regulation by Foxp3-ΔE2 Tregs and reduced Foxp3-ΔE2 Treg localization to the T cell zone of the lung-draining lymph node during T cell priming contributed to defective CD8^+^ T cell suppression. These data support a model where dysfunctional programming in Foxp3-ΔE2 effector Tregs promotes CD8^+^ T-cell responses during infection.

## RESULTS

### Dysfunctional effector programming in Foxp3-ΔE2 Tregs

Initial studies in Foxp3-ΔE2 mice revealed altered ratios of central and effector Tregs in Foxp3-ΔE2 mice and identified several hundred differentially expressed genes (DEGs) between total Foxp3-ΔE2 and Foxp3-FL Tregs (*15*). These data led us to question whether the DEGs between Foxp3-FL and Foxp3-ΔE2 Tregs were truly driven by isoform-specific differences in transcriptional programming or instead simply reflected increased central Treg skewing in Foxp3-ΔE2 Tregs. To address this question, we generated dual reporter mice that allow live sorting of Foxp3-FL (eGFP^+^ mRFP^+^) and Foxp3-ΔE2 (eGFP^+^ mRFP^-^) Tregs based on fluorescent reporter expression (Fig 1A). We sorted splenic central (CD62L^hi^ CD44^lo^) or effector (CD44^hi^ CD62L^lo^) Foxp3-FL and Foxp3-ΔE2 Tregs (Fig 1B) and total splenic Foxp3-FL and Foxp3-ΔE2 Tregs (Supp Fig 1A) from these mice then performed transcriptomic analyses by bulk RNA-seq.

**Fig. 1.**
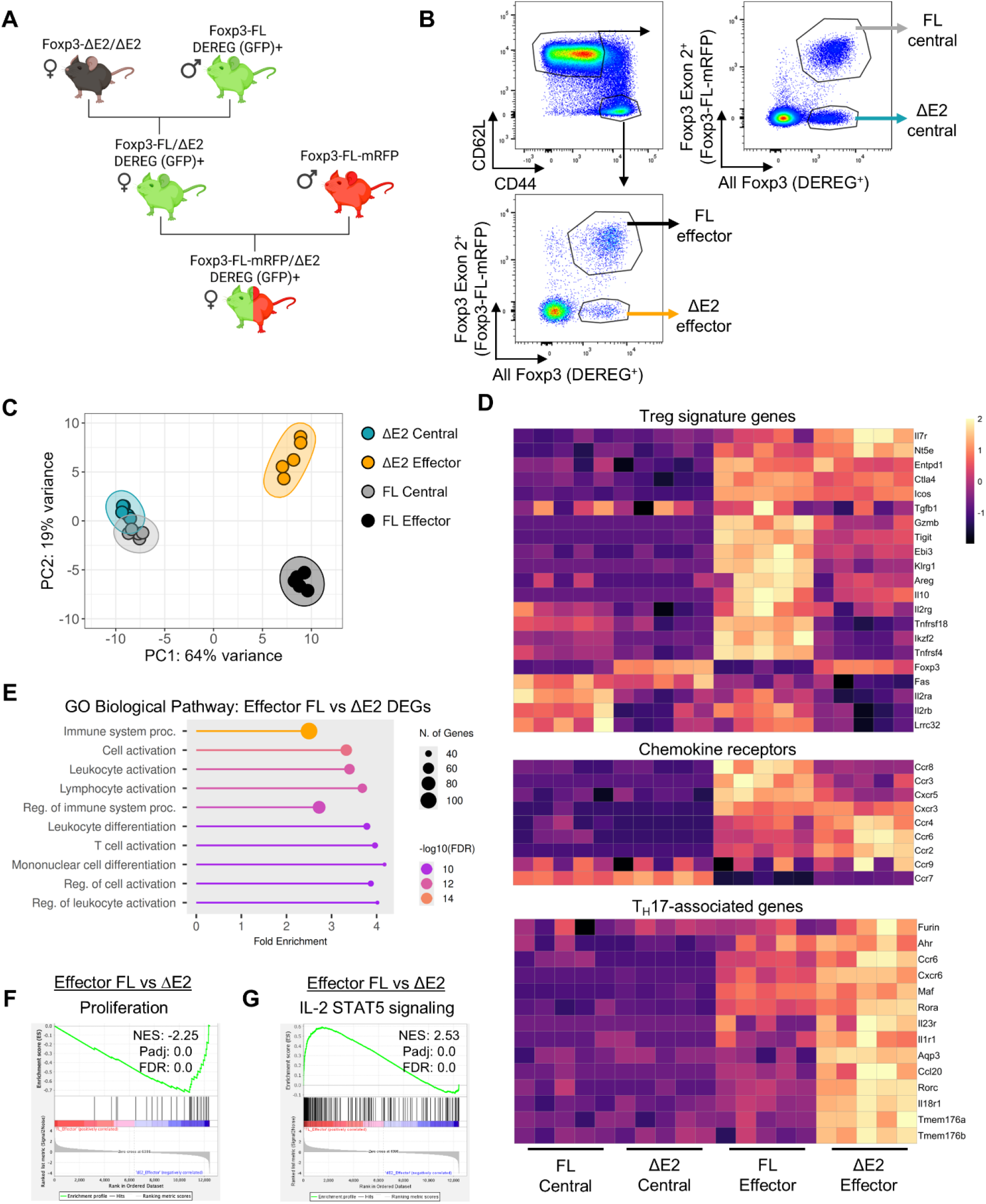
Dysfunctional effector programming in Foxp3-ΔE2 Tregs. **(A)** Dual reporter Foxp3-FL-mRFP/ΔE2 DEREG^+^ mice were generated to allow separation of live Foxp3-ΔE2 and Foxp3-FL Tregs. **(B)** A representative flow plot of central (CD62L^hi^ CD44^lo^) and effector (CD44^hi^ CD62L^lo^) Foxp3-ΔE2 and Foxp3-FL Tregs from dual reporter mice which were sorted then analyzed by bulk RNA-seq. **(C)** A PCA plot with 95% confidence ellipses of central and effector Foxp3-FL and Foxp3-ΔE2 Tregs. **(D)** A Z-scored heatmap of Treg signature genes, differentially expressed chemokine receptors, and T_H_17-associated genes. **(E)** A GO Biological Pathway analysis of effector Foxp3-FL versus Foxp3-ΔE2 Treg DEGs. **(F)** GSEA of effector Foxp3-FL versus Foxp3-ΔE2 Tregs for a proliferation gene set and **(G)** hallmark IL-2 STAT5 signaling genes with normalized enrichment score (NES), adjusted p-value (padj), and false-discovery rate (FDR) shown. *n*=4 mice.

We found 144 DEGs between central Foxp3-FL and Foxp3-ΔE2 Tregs, 431 DEGs between effector Foxp3-FL and Foxp3-ΔE2 Tregs, and 700 DEGs between total Foxp3-FL and Foxp3-ΔE2 Tregs (Supp Table 1-3, Supp Fig 1B-D). To identify which DEGs amongst total Foxp3-ΔE2 Tregs were driven by central Treg skewing, we overlapped central versus effector differentiating genes with total Foxp3-ΔE2 versus Foxp3-FL DEGs and found 124 genes driven by central versus effector Treg programming (Supp Fig 2A-B), including *Bcl2*, *Bach2*, *Irf4*, *Itgae*, *Batf*, and *Bhlhe40*. Thus, many previously identified DEGs in total Foxp3-ΔE2 Tregs may be driven by altered central versus effector Treg differentiation and may not be directly regulated by Foxp3 Exon 2.

**Fig. 2.**
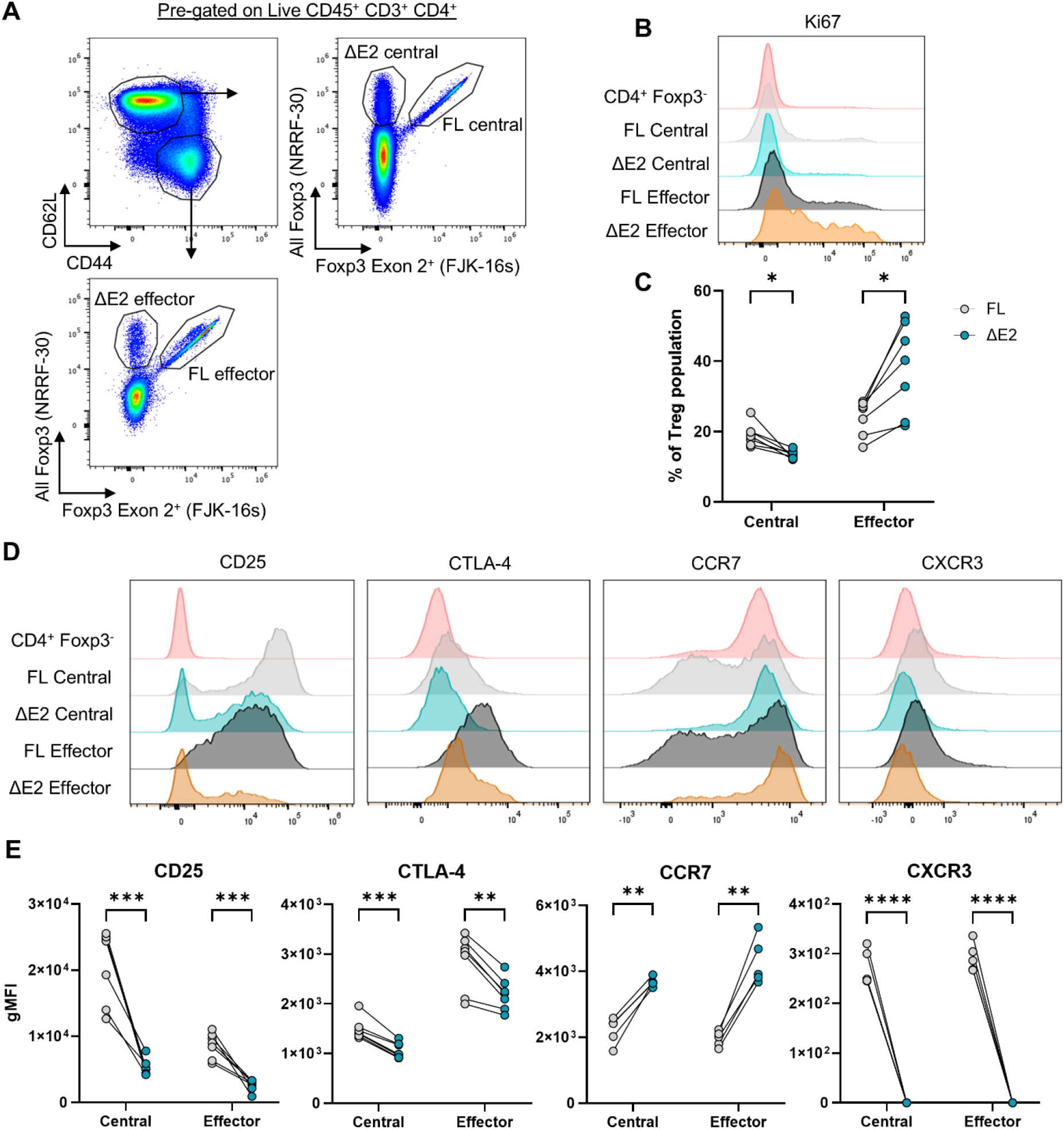
Altered expression of key regulatory and homing molecules in central and effector Foxp3-ΔE2 Tregs. **(A)** Gating strategy used to identify central and effector Foxp3-FL and Foxp3-ΔE2 Tregs from homeostatic Foxp3-FL/ΔE2 female mice aged 8-12 weeks. **(B)** Representative flow histogram of Ki67 expression on central and effector Foxp3-FL and Foxp3-ΔE2 Tregs with CD4^+^ Foxp3^-^cells shown as an internal control and **(C)** percentage of Ki67^+^ central and effector Foxp3-FL and Foxp3-ΔE2 Tregs. **(D)** Representative flow histograms of CD25, CTLA-4, CCR7, and CXCR3 expression on central and effector Foxp3-FL and Foxp3-ΔE2 Tregs with CD4^+^ Foxp3^-^cells shown as an internal control. **(E)** Geometric mean fluorescence intensity (gMFI) of CD25, CTLA-4, CCR7, and CXCR3 on central and effector Foxp3-FL and Foxp3-ΔE2 Tregs. *n*=6 mice. Pairing is shown within each mouse. \**p*<0.05, \*\**p*<0.01, \*\*\**p*<0.001, \*\*\*\**p*<0.0001 by two-tailed paired t tests with Holm-Sidak method of correction for multiple comparisons.

We next analyzed isoform-specific differences in transcriptional programming between Foxp3-ΔE2 and Foxp3-FL Tregs within central and effector populations. By principal component analysis (PCA), the differences between Treg populations were largely driven by differentiation state in PC1 and genotype in PC2 with a far greater magnitude of transcriptional difference found between Foxp3-FL and Foxp3-ΔE2 effector Tregs (Fig 1C). Top variance-driving DEGs between the four populations of central and effector Foxp3-FL and Foxp3-ΔE2 Tregs included Treg signature genes *Il2ra*, *Il2rb*, and *Tnfrsf18*; chemokine receptors that control Treg localization; and T_H_17-associated genes *Rorc*, *Cxcr6*, and *Tmem176b* (Supp Fig 3A). Given the importance of Foxp3 in driving Treg transcriptional programming and the known role of Foxp3 Exon 2 in antagonizing T_H_17 programming (*13*), we further compared expression levels of an expanded list of Treg signature genes, differentially expressed chemokine receptors, and T_H_17-signature genes (Fig 1D). Treg signature genes were overall downregulated in Foxp3-ΔE2 Tregs, especially in the effector Treg population. Similarly, the strongest differences in chemokine receptor expression were between Foxp3-FL and Foxp3-ΔE2 effector Tregs. Most strikingly, T_H_17-associated genes were substantially upregulated in Foxp3-ΔE2 effector Tregs but not in central or total Foxp3-ΔE2 Tregs (Fig 1D, Supp Fig 3B).

**Fig. 3.**
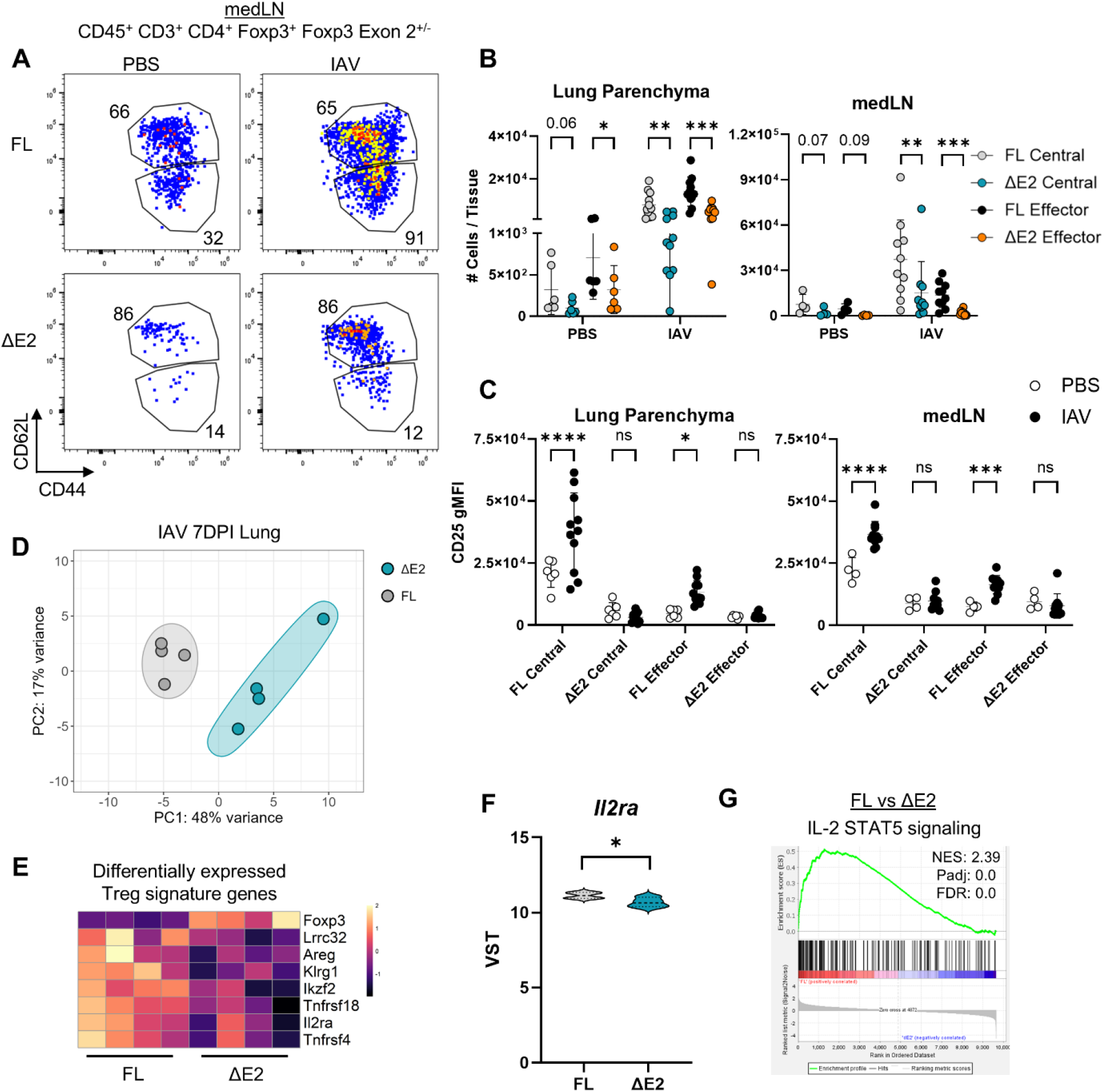
Exacerbated Foxp3-ΔE2 effector Treg dysfunction in influenza viral infection. Foxp3-FL/ΔE2 mice aged 8-12 weeks were infected with X31 IAV then analyzed during acute infection at 7DPI. **(A)** Representative flow plots of medLN central and effector Foxp3-FL and Foxp3-ΔE2 Treg populations in PBS mock infected or IAV infected mice. **(B)** Number of central and effector Foxp3-FL and Foxp3-ΔE2 Tregs in the lung parenchyma or the lung-draining mediastinal lymph node (medLN). **(C)** CD25 gMFI expression on central and effector Foxp3-FL and Foxp3-ΔE2 Tregs in the lung parenchyma and medLN. **(D-G)** Foxp3-FL and Foxp3-ΔE2 Tregs were sorted from the lung parenchyma of X31 IAV infected dual reporter Foxp3-FL-mRFP/ΔE2 DEREG^+^ mice at 7DPI then analyzed by bulk RNA-seq. **(D)** A PCA plot with 95% confidence ellipses of Foxp3-FL and Foxp3-ΔE2 Tregs. **(E)** A Z-scored heatmap of differentially expressed Treg signature genes between Foxp3-FL and Foxp3-ΔE2 Tregs. **(F)** Variance stabilizing transformed (VST) *Il2ra* expression in Foxp3-FL and Foxp3-ΔE2 Tregs. **(G)** GSEA of hallmark IL-2/STAT5 signaling genes in Foxp3-FL vs Foxp3-ΔE2 Tregs with NES, padj, and FDR shown. *n*=4-9 mice per group. Bars show mean values; error bars show SD. \**p*<0.05, \*\**p*<0.01, \*\*\**p*<0.001, \*\*\*\**p*<0.0001 by two-tailed paired t test with Holm-Sidak method of correction for multiple comparisons within central or effector Treg populations (B), unpaired two-tailed t tests with Holm-Sidak method of correction for multiple comparisons (C), or Wald Test performed within DESeq2 with adjusted p-value reported (F).

GO Biological pathway analysis of DEGs between Foxp3-FL versus Foxp3-ΔE2 Tregs identified an enrichment in T cell activation related pathways (Fig 1E, Supp Fig 4A). Amongst T cell activation genes, *Mki67* expression was significantly increased in Foxp3-ΔE2 effector Tregs (Supp Fig 4B) which corresponded to an enrichment for proliferation signature genes in effector Foxp3-ΔE2 Tregs by a gene set enrichment analysis (GSEA, Fig 1F, Supp Fig 4C). Finally, the most significantly enriched hallmark gene set for both central and effector Foxp3-FL versus Foxp3-ΔE2 Tregs was IL-2/STAT5 signaling genes (Fig 1G, Supp Fig 4D).

**Fig. 4.**
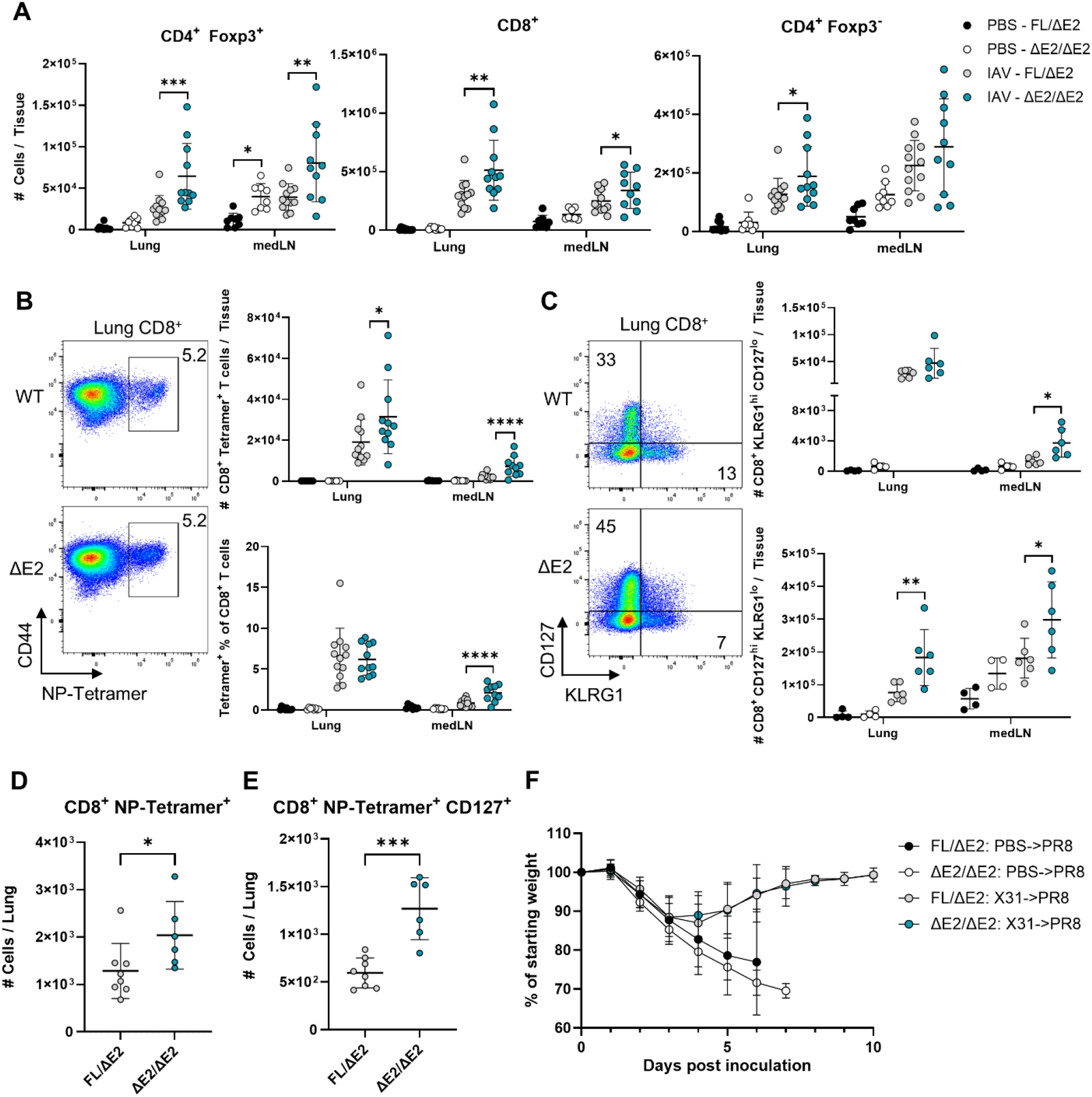
Enhanced magnitude of CD8^+^ T cell responses during influena viral infection in Foxp3-ΔE2/ΔE2 mice. **(A-C)** Foxp3-FL/ΔE2 and Foxp3-ΔE2/ΔE2 mice aged 8-12 weeks were PBS mock infected or infected with X31 IAV then analyzed during acute infection at 7DPI. **(A)** Numbers of CD4^+^ Foxp3^+^, CD8^+^, and CD4^+^ Foxp3^-^T cells in the lung parenchyma and medLN. **(B)** Representative flow plots, counts, and percentages of CD8^+^ NP-Tetramer^+^ T cells in the lung parenchyma and medLN. **(C)** Representative flow plots and counts of CD8^+^ KLRG1^hi^ CD127^lo^ and CD8^+^ CD127^hi^ KLRG1^lo^ T cells in the lung parenchyma and medLN. **(D-E)** Foxp3-FL/ΔE2 and Foxp3-ΔE2/ΔE2 mice were infected with X31 IAV and analyzed at 30DPI. **(D)** Number of CD8^+^ NP-Tetramer^+^ T cells in the lung parenchyma. **(E)** Number of CD8^+^ NP-Tetramer^+^ CD127^+^ T cells in the lung parenchyma. **(F)** Foxp3-FL/ΔE2 or Foxp3-ΔE2/ΔE2 mice were PBS mock infected or X31 IAV infected, recovered for 30 days, then inoculated with a lethal dose of PR8 IAV. Weight loss is shown as a percentage of each animal’s starting weight. *n*=4-11 mice per group. Bars show mean values; error bars show SD. \**p*<0.05, \*\**p*<0.01, \*\*\**p*<0.001,\*\*\*\**p*<0.0001 by two-way ANOVA followed by post-hoc t tests (A-C), two-tailed unpaired t tests with Holm-Sidak method of correction for multiple comparisons (D-E), and generalized additive mixed model (F).

Confirming our transcriptional analysis, we found an increased percentage of Ki67^+^ cells amongst effector Foxp3-ΔE2 Tregs (Fig 2A-C); decreased CD25, CTLA-4, and ICOS expression on Foxp3-ΔE2 central and effector Tregs; and increased CD40L and PD-1 expression on Foxp3-ΔE2 central and effector Tregs (Fig 2D-E, Supp Fig 5A-B). CCR7 expression was increased on Foxp3-ΔE2 central and effector Tregs, and CCR4, CCR8, CXCR3, and CXCR5 expression was decreased (Fig 2D-E, Supp Fig 5C). Taken together, these data identify key transcriptional and phenotypic differences in central and effector Foxp3-ΔE2 Tregs. Additionally, by parsing apart central and effector populations, we identified that Foxp3 Exon 2 plays a more pronounced role in effector Tregs by maintaining expression of Treg signature genes, inhibiting T_H_17-like differentiation, and driving chemokine receptor expression required for proper cellular localization.

**Fig. 5.**
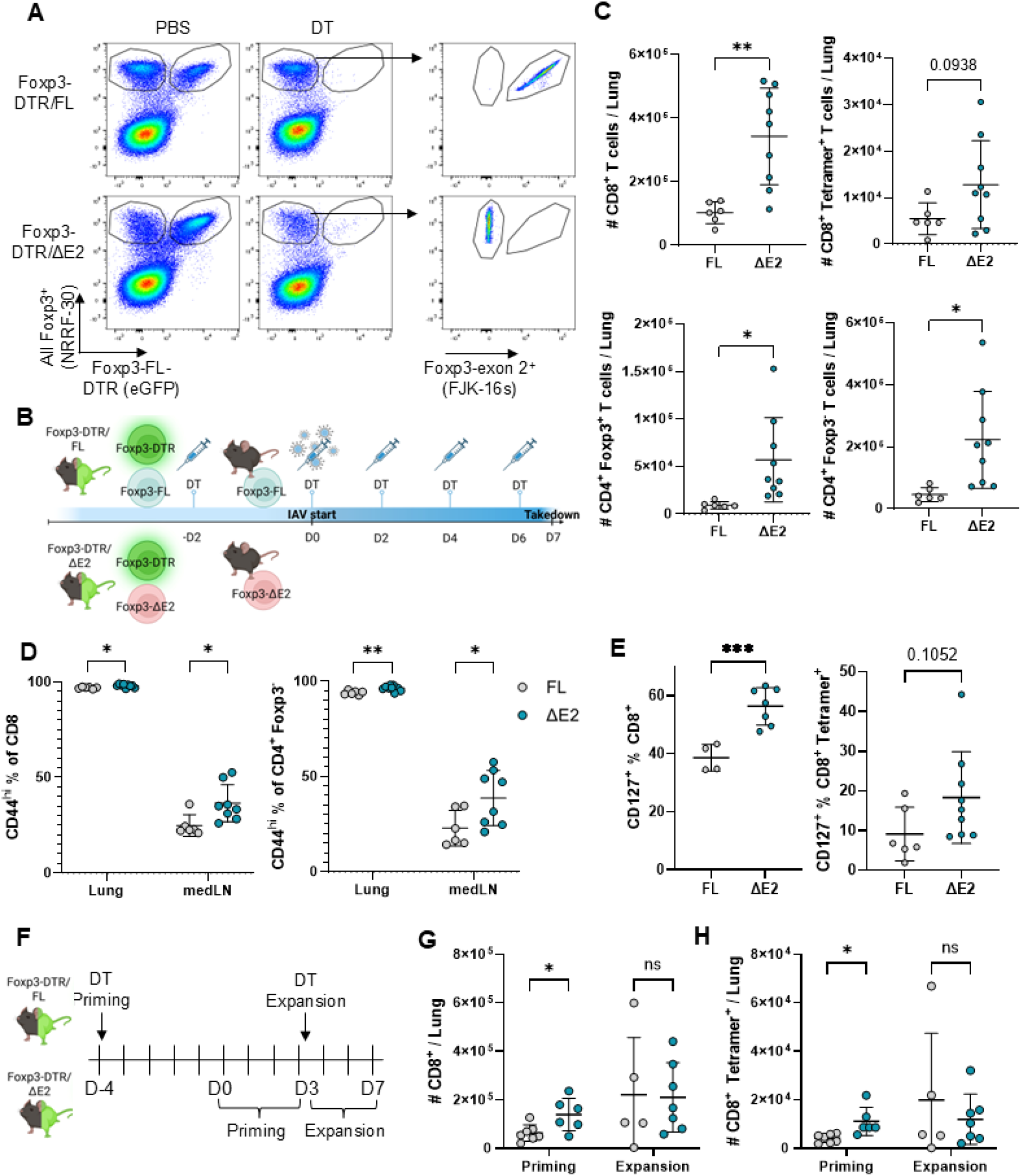
Defects in CD8^+^ T cell regulation by Foxp3-ΔE2 Tregs occur during priming. Foxp3-DTR/FL and Foxp3-DTR/ΔE2 mice were bred to allow generation of Foxp3-FL or Foxp3-ΔE2 mice upon DT treatment with no prior systemic inflammation. **(A)** Representative flow plots showing the efficacy of Foxp3-FL and Foxp3-ΔE2 mouse generation following treatment with 50µg/kg body weight DT once two days prior to analysis. **(B)** Experiment schematic for X31 IAV infection in Foxp3-DTR x Foxp3-ΔE2 mice. **(C-E)** Foxp3-ΔE2 or Foxp3-FL mice were generated via DT treatment two days prior to infection with X31 IAV then maintained with DT treatment every other day. Mice were analyzed at 7DPI with X31 IAV. **(C)** Numbers of CD8^+^ T cells, CD8^+^ NP-Tetramer^+^ T cells, CD4^+^ Foxp3^+^, and CD4^+^ Foxp3^-^T cells in the lung parenchyma. **(D)** Percentages of CD44^hi^ CD8^+^ and CD44^hi^ CD4^+^ Foxp3^-^T cells in the lung parenchyma and medLN. **(E)** Percentages of CD8^+^ CD127^+^ and CD8^+^ NP-Tetramer^+^ CD127^+^ T cells in the lung parenchyma. **(F-H)** Foxp3-DTR x Foxp3-ΔE2 mice were DT treated either at -4DPI or +3DPI with X31 IAV to determine if Foxp3-ΔE2 Tregs fail to restrain CD8^+^ T cells during priming (−4DPI DT treatment) or expansion (+3DPI DT treatment). **(F)** Experiment schematic. **(G)** Number of CD8^+^ and **(H)** CD8^+^ NP-Tetramer^+^ T cells in the lung parenchyma at 7DPI. *n*=5-9 mice per group. Bars show mean values; error bars show SD. \**p*<0.05, \*\**p*<0.01, \*\*\**p*<0.001, \*\*\*\**p*<0.0001 by two-tailed unpaired t tests (C & E) and two-way ANOVA followed by post-hoc t tests (D & G-H).

### Exacerbated Foxp3-ΔE2 effector Treg dysfunction during inflammation

To determine how Foxp3-ΔE2 central and effector Tregs respond to inflammation, we used an influenza A viral (IAV) infection model. Foxp3-FL/ΔE2 heterozygous female mice were either PBS mock infected or IAV infected then analyzed during the acute phase of infection at 7 days post inoculation (DPI). We found a significantly lower number of total Foxp3-ΔE2 than Foxp3-FL Tregs in the lung parenchyma and mediastinal lymph node (medLN) in IAV infected mice (Supp Fig 6A). Similarly, although the total number of cells increased during infection, there were fewer central and effector Foxp3-ΔE2 than Foxp3-FL Tregs in both PBS treated and IAV infected mice (Fig 3A-B, Supp Fig 6B-C). Notably, CD25 expression increased on Foxp3-FL central and effector Tregs during infection but failed to upregulate on Foxp3-ΔE2 central Tregs and decreased on medLN Foxp3-ΔE2 effector Tregs, suggesting diminished regulation of IL-2 availability by Foxp3-ΔE2 Tregs during inflammation (Fig 3C).

**Fig. 6.**
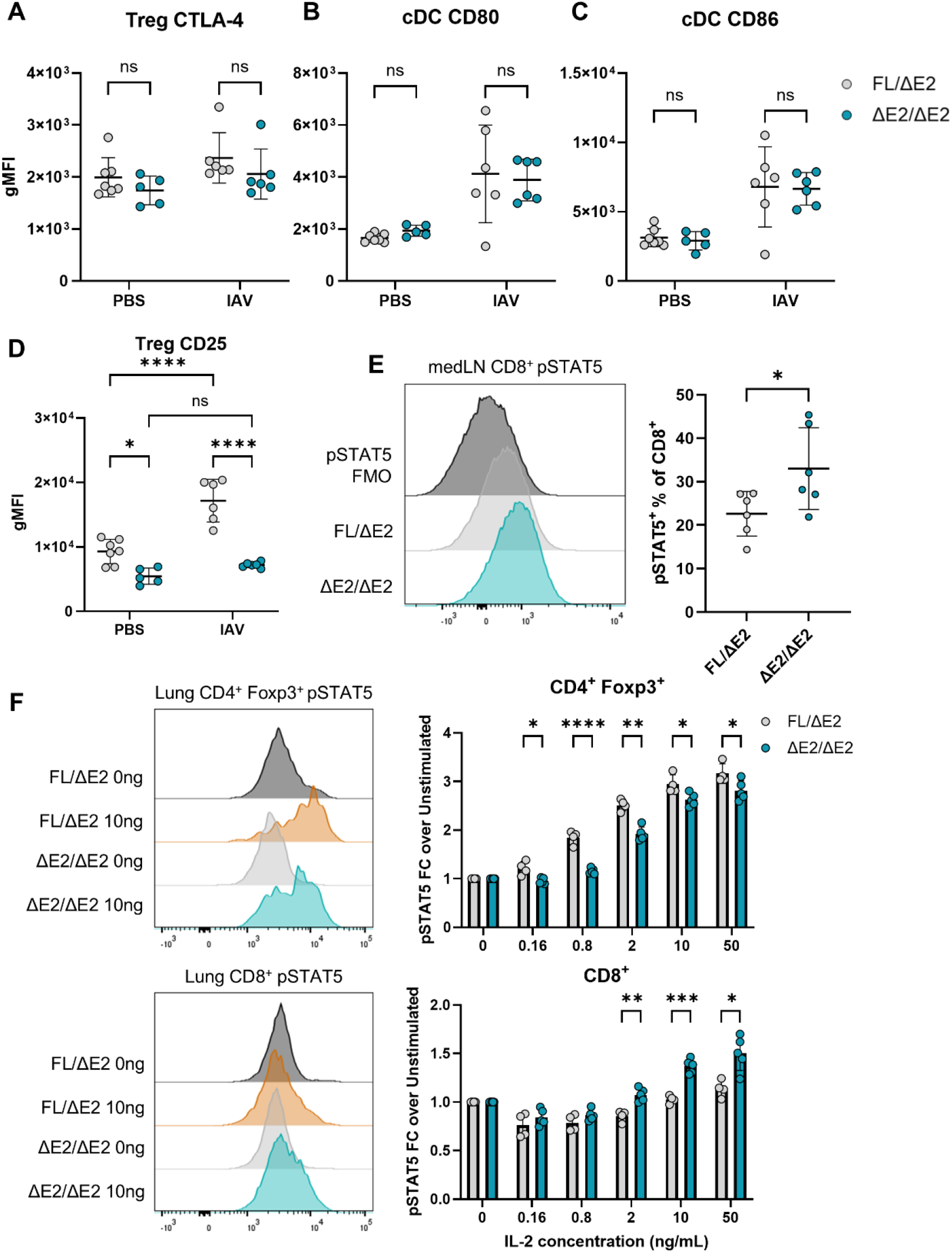
Reduced IL-2 sink activity by Foxp3-ΔE2 Tregs associates with increased IL-2 signaling in CD8^+^ T cells during priming. **(A-E)** Foxp3-FL/ΔE2 and Foxp3-ΔE2/ΔE2 mice were PBS mock infected or infected with X31 IAV then analyzed during CD8^+^ T cell priming at 3DPI in the medLN. gMFI expression of **(A)** CTLA-4 on Tregs, **(B)** CD80 on classical dendritic cells (cDCs), **(C)** CD86 on cDCs, and **(D)** CD25 on Tregs. **(E)** A representative histogram of pSTAT5 levels on CD8^+^ T cells from the medLN directly *ex vivo* and the percentage of pSTAT5^+^ cells amongst total CD8^+^ T cells. **(F)** Foxp3-FL/ΔE2 and Foxp3-ΔE2/ΔE2 mice were infected with X31 IAV. At 7DPI, lung cells were stimulated with indicated concentrations of IL-2 *ex vivo* then analyzed for pSTAT5 induction in CD4^+^ Foxp3^+^ and CD8^+^ T cells by flow cytometry. Representative histograms of pSTAT5 staining and the pSTAT5 fold change over unstimulated cells within each mouse are shown. *n*=5-6 mice per group. Bars show mean values; error bars show SD. \**p*<0.05, \*\**p*<0.01, \*\*\**p*<0.001, \*\*\*\**p*<0.0001 by two-way ANOVA followed by multiple t tests (A-D) or unpaired t tests with Holm-Sidak method of correction for multiple comparisons as appropriate (E-F).

Since the majority of lung parenchymal Tregs during IAV infection are effector differentiated (Supp Fig 6B-C), we sorted lung parenchymal Foxp3-ΔE2 and Foxp3-FL Tregs from dual reporter Foxp3-FL-mRFP/ΔE2 DEREG^+^ mice for transcriptomic analysis. We found that Treg populations separated by genotype in a PCA (Fig 3D) and there were 457 DEGs between Foxp3-ΔE2 and Foxp3-FL Tregs (Supp Table 4). Among these DEGs, 65 overlapped with steady-state effector Foxp3-FL vs Foxp3-ΔE2 Treg DEGs, including *Tcf7*, *Lrrc32*, *Ikzf2*, and *Ikzf4* (Supp Fig 6D). Other DEGs of interest included Treg signature genes such as *Tnfrsf18*, *Areg*, and *Il2ra* (Fig 3E-F, Supp Fig 6E). As in the steady-state, GSEA identified the most significantly enriched hallmark gene set in Foxp3-FL versus Foxp3-ΔE2 Tregs during infection was IL-2/STAT5 signaling genes (Fig 3G). Taken together, these data demonstrate that inflammatory challenge amplifies Foxp3-ΔE2 effector Treg dysfunction, highlighting a critical role for Foxp3 Exon 2 in upregulating CD25 expression and controlling IL-2 responsiveness during inflammation.

### Increased magnitude of CD8^+^ T cell response in influenza infected Foxp3-ΔE2 mice

We next set out to determine how exacerbated Foxp3-ΔE2 effector Treg dysfunction during inflammation affects conventional T cell regulation. Foxp3-FL/ΔE2 mice have a comparable immune response to influenza infection as Foxp3-FL/FL mice (Supp Fig 7A-B). Therefore, we used homozygous Foxp3-ΔE2/ΔE2 mice to identify the effects of Foxp3-ΔE2 Tregs on conventional T cells during infection. At 7DPI with X31 IAV, total CD4^+^ Foxp3^+^ and CD8^+^ T cell numbers were substantially increased in the lung parenchyma and medLN of Foxp3-ΔE2/ΔE2 mice while CD4^+^ Foxp3^-^T cell numbers were only modestly increased (Fig 4A). Additionally, an increased percentage of CD8^+^ T cells in Foxp3-ΔE2/ΔE2 mice were activated and expressing Granzyme B in the medLN during infection (Supp Fig 8A-B).

**Fig. 7.**
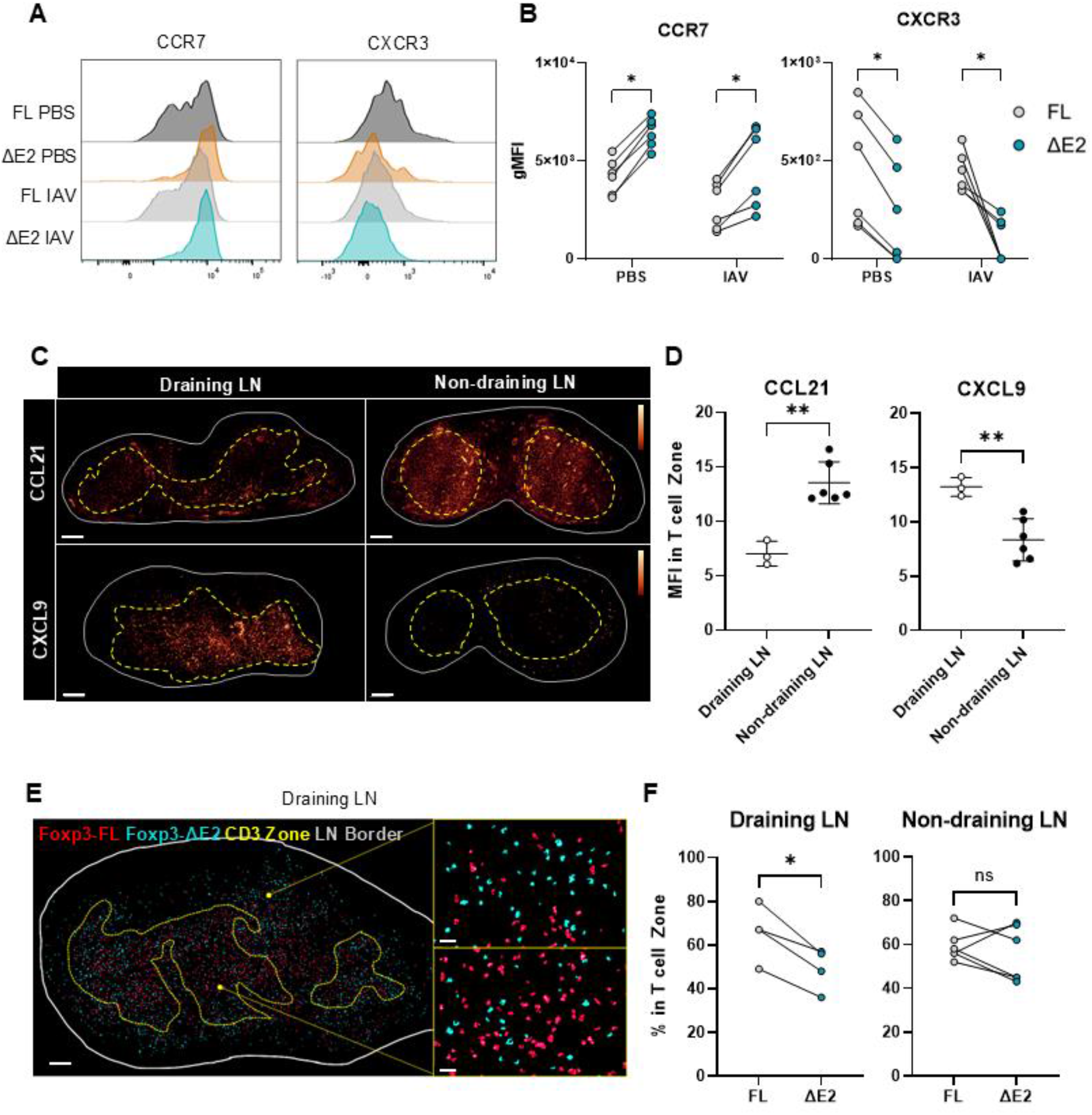
Foxp3-ΔE2 Tregs have reduced localization to the T cell zone in the lung-draining lymph node during priming. **(A-B)** Foxp3-FL/ΔE2 mice were PBS mock infected or X31 IAV infected and chemokine receptor expression levels were measured on Foxp3-FL and Foxp3-ΔE2 Tregs at 3DPI in the medLN by flow cytometry. **(A)** Representative histograms and **(B)** gMFI of CCR7 and CXCR3. **(C-F)** Dual reporter Foxp3-FL-mRFP/ΔE2 DEREG^+^ mice were infected with X31 IAV then analyzed at 3DPI by confocal microscopy. **(C)** Representative images of CCL21 and CXCL9 staining in the draining and non-draining LN (scale bar=200μm) with LUT bars shown. **(D)** CCL21 and CXCL9 median fluorescence intensity (MFI) in the draining and non-draining LN. **(E)** Representative images of Foxp3-FL (red) and Foxp3-ΔE2 (cyan) Treg localization within and outside of the T cell zone (yellow dashed line) of the draining LN (scale bar=150μm for whole LN and 20μm for zoom-in). **(F)** The percentage of Foxp3-FL or Foxp3-ΔE2 Tregs in the T cell zone in the draining or non-draining LNs. *n*=3-6 mice. \**p*<0.05, \*\**p*<0.01 by two-tailed paired t tests with Holm-Sidak method of correction for multiple comparisons as appropriate (B & F) or two-tailed unpaired t tests (D).

Given the established function of Tregs in restraining CD8^+^ T cell responses through limiting IL-2 availability (*6*), we subsequently focused our analyses on CD8^+^ T cell responses in IAV-infected mice. We used an influenza nucleoprotein (NP) specific T cell tetramer to study the antigen-specific CD8^+^ T cell response and found an increased number of CD8^+^ NP-Tetramer^+^ cells in the lungs and medLN in Foxp3-ΔE2/ΔE2 mice and an increased percentage of NP-Tetramer^+^ cells amongst CD8^+^ T cells in Foxp3-ΔE2/ΔE2 medLNs (Fig 4B). Despite the increased number of activated and cytolytic CD8^+^ T cells in Foxp3-ΔE2/ΔE2 mice, we observed no difference in weight loss (Supp Fig 8C), lung pathology (Supp Fig 8D-E), or viral mRNA levels (Supp Fig 8F).

We found an increased number of CD8^+^ CD127^hi^ KLRG1^lo^ memory precursors, but not CD8^+^ KLRG1^hi^ CD127^lo^ short-lived effectors, in infected Foxp3-ΔE2/ΔE2 mice (Fig 4C, Supp Fig 8G). At an early memory timepoint (30DPI), the total number of CD8^+^ T cells in the lungs had contracted (Supp Fig 9A) but there were an increased number of CD8^+^ NP-Tetramer^+^ T cells and CD127^+^ NP-Tetramer^+^ T cells in the lung parenchyma of Foxp3-ΔE2/ΔE2 mice (Fig 4D-E). This corresponded to enhanced long-term persistence of CD8^+^ NP-Tetramer^+^ cells in Foxp3-ΔE2/ΔE2 mice at 90DPI (Supp Fig 9B). To measure memory CD8^+^ T cell function in Foxp3-ΔE2/ΔE2 mice, we used a heterosubtypic infection model where mice that were PBS mock infected or X31 IAV infected 30 days previously were re-infected with the serologically distinct PR8 IAV strain. We observed no difference in weight loss, survival, or CD8^+^ T cell counts between Foxp3-ΔE2/ΔE2 and Foxp3-FL/ΔE2 mice (Fig 4F & Supp Fig 9C-D), demonstrating a functional memory CD8^+^ T cell compartment in Foxp3-ΔE2/ΔE2 mice.

### Cell-intrinsic defects in Foxp3-ΔE2 Tregs drive enhanced CD8^+^ T cell responses

To determine whether the diminished restraint of CD8^+^ T cell responses in Foxp3-ΔE2 mice reflects an intrinsic defect in Foxp3-ΔE2 Treg function rather than the chronic inflammatory environment in these mice, we developed a system to acutely replace wild-type Tregs with Foxp3-ΔE2 Tregs. We crossed Foxp3-DTR male mice to female Foxp3-FL/ΔE2 mice to generate Foxp3-DTR/FL and Foxp3-DTR/ΔE2 female offspring (Supp Fig 10A). These mice have no differences in inflammation at steady-state (Supp Fig 10B-D) although, similar to Foxp3-FL/ΔE2 mice, Foxp3-DTR Tregs outcompete Foxp3-ΔE2 Tregs in Foxp3-DTR/ΔE2 mice. Administering diphtheria toxin (DT) to these mice depletes Foxp3-FL-expressing Foxp3-DTR Tregs, generating Foxp3-FL or Foxp3-ΔE2 mice without prior differences in inflammation (Fig 5A). We DT treated Foxp3-DTR x Foxp3-ΔE2 mice, infected with IAV two days later, and DT treated every other day to maintain depletion until analysis at 7DPI (Fig 5B). We found significantly increased weight loss in Foxp3-ΔE2 mice and a trend toward decreased viral RNA levels in Foxp3-ΔE2 mouse lungs (Supp Fig 10E-F). We also found an increased number and activation of T cells in Foxp3-ΔE2 mice (Fig 5C-D) as well as increased memory precursor differentiation (Fig 5E). Thus, defects in CD8^+^ T cell restraint are due to intrinsic Treg programming defects in Foxp3-ΔE2 Tregs, not simply due to underlying systemic inflammation in Foxp3-ΔE2 mice.

To determine when Foxp3-ΔE2 Tregs are deficient in CD8^+^ T cell regulation, we either DT depleted Foxp3-DTR Tregs at 4 days prior to infection to eliminate WT Tregs during the initial priming phase (Foxp3-DTR Tregs replete at about 7 days post DT treatment) or at 3 days after the start of infection for the expansion phase (Fig 5F). We found that total CD8^+^ and CD8^+^ NP-Tetramer^+^ T cells were increased in Foxp3-ΔE2 mice DT treated during priming but not expansion (Fig 5G-H), indicating that the defect in CD8^+^ T cell suppression by Foxp3-ΔE2 Tregs occurs during priming.

### Diminished IL-2R-mediated regulation of CD8^+^ T cells by Foxp3-ΔE2 Tregs

To identify potential mechanisms by which Foxp3-ΔE2 Tregs fail to restrain CD8^+^ T cell responses, we evaluated CTLA-4-dependent transendocytosis of costimulatory ligands and IL-2R-mediated sequestration of IL-2 by Foxp3-ΔE2 Tregs. We selected these pathways because they have a well-described role in Treg regulation of conventional T cells, and Foxp3-ΔE2 effector Tregs have a cell-intrinsic reduction in the expression of CTLA-4 and CD25 at steady-state.

To assess CTLA-4 function in Foxp3-ΔE2 Tregs, we first performed an *in vitro* transendocytosis assay. MACS-enriched CD4^+^ T cells from Foxp3-ΔE2/ΔE2 or Foxp3-FL/ΔE2 mice were co-cultured with CD86-eGFP expressing 3T3 cells, and we quantified CTLA-4-dependent CD86-eGFP uptake by Tregs (Supp Fig 11A-B) (*22*). Foxp3-ΔE2 Tregs from heterozygous female Foxp3-FL/ΔE2 mice, but not homozygous Foxp3-ΔE2/ΔE2 mice, had reduced CD86 transendocytosis, consistent with reduced CTLA-4 expression on Foxp3-ΔE2 Tregs within heterozygous mice only (Supp Fig 11C-D) (*15*). To define the role of CTLA-4 in Foxp3-ΔE2 Treg suppression of CD8^+^ T cells *in vivo*, we infected Foxp3-ΔE2/ΔE2 or Foxp3-FL/ΔE2 mice with IAV and analyzed the medLN during priming at 3DPI. We found no difference in the expression levels of CTLA-4 on Tregs (Fig 6A) or CD80 and CD86 on classical dendritic cells (cDCs, Fig 6B-C), indicating that CTLA-4 regulation of co-stimulatory molecules by Foxp3-ΔE2 Tregs was intact during T cell priming *in vivo*.

Next, to measure the ability of Foxp3-ΔE2 Tregs to respond to IL-2, we stimulated MACS-enriched CD4^+^ T cells from Foxp3-ΔE2 or Foxp3-FL mice with IL-2 then measured pSTAT5 induction (Supp Fig 11E). pSTAT5 induction was lower in Foxp3-ΔE2 Tregs across several concentrations of IL-2, indicating that Foxp3-ΔE2 Tregs have a reduced capacity to signal in response to IL-2 (Supp Fig 11F). During IAV infection, we found that CD25 expression was significantly lower on Foxp3-ΔE2 Tregs and failed to upregulate with infection at both 3DPI and 7DPI (Fig 6D, Supp Fig 12A-E). Reciprocally, CD25 expression was modestly increased on CD8^+^ T cells in newly generated Foxp3-ΔE2 mice at 7DPI with IAV (Supp Fig 12E). Together, these data strongly suggested a diminished ability of Foxp3-ΔE2 Tregs to regulate IL-2 levels *in vivo*.

A significantly increased percentage of CD8^+^ T cells assessed directly *ex vivo* at 3DPI with IAV were pSTAT5^+^ in Foxp3-ΔE2 mice (Fig 6E, Supp Fig 12F), indicating increased IL-2 signaling in CD8^+^ T cells from Foxp3-ΔE2 mice during priming. Furthermore, we observed decreased pSTAT5 induction in CD4^+^ Foxp3^+^ Tregs and increased pSTAT5 induction in CD8^+^ T cells in response to IL-2 stimulation *ex vivo* (Fig 6F). These data indicate that impaired IL-2 regulation by Foxp3-ΔE2 Tregs during priming plays a role in their deficient regulation of CD8^+^ T cells.

### Reduced Foxp3-ΔE2 Treg localization to the lymph node T cell zone during priming

In addition to impaired CD25 expression, Foxp3-ΔE2 Tregs exhibited altered expression levels of tissue-homing chemokine receptors at steady-state, including CCR7 and CXCR3 (Fig 2D-E). CXCR3 expressing Tregs play a key role in CD8^+^ T cell regulation in tumor models (*23, 24*).

Therefore, altered chemokine receptor expression on Foxp3-ΔE2 Tregs may change cellular positioning, contributing to aberrant regulation of CD8^+^ T cell responses during IAV infection. Since Foxp3-ΔE2 Tregs failed to regulate CD8^+^ T cell responses during the priming phase of infection (Fig 5G-H), we measured chemokine receptor expression on Foxp3-ΔE2 Tregs in the medLN at 3DPI with X31 IAV. Similar to steady-state, Foxp3-ΔE2 Tregs had increased CCR7 expression and decreased CCR4, CCR5, CXCR3, and CXCR5 expression during infection (Fig 7A-B, Supp Fig 13A), suggesting altered Foxp3-ΔE2 Treg localization in the lung-draining lymph node during infection.

Chemokine expression and Treg localization were quantified in IAV infected dual reporter Foxp3-FL-mRFP/ΔE2 DEREG^+^ mice at 3DPI by confocal microscopy. We found decreased CCL21 expression and increased CXCL9 expression in the lung-draining medLN T cell zone compared with the non-draining (inguinal) LN (Fig 7C–D), similar to reports in other settings of inflammation (*25–29*), suggesting that Foxp3-ΔE2 Tregs may have reduced recruitment to the T cell zone. We identified Foxp3-FL (mRFP^+^ eGFP^+^) and Foxp3-ΔE2 (mRFP^-^eGFP^+^) Treg populations through fluorescent reporter expression (Supp Fig 13B-C) and quantified their localization within the T cell zone (Fig 7E, Supp Fig 13D). In the draining LN, a significantly lower percentage of Foxp3-ΔE2 Tregs localized to the T cell zone whereas no such positional difference was observed in the non-draining LN (Fig 7F, Supp Fig 13E). Taken together, altered expression of chemokine receptors on Foxp3-ΔE2 Tregs drives reduced co-localization with conventional T cells in the inflamed lymph node during priming, contributing to reduced suppression of CD8^+^ T cells.

## DISCUSSION

Tregs are a heterogeneous immune cell subset with central and effector populations. In mice, central Tregs are CD44^lo^ CD62L^hi^ CCR7^+^ and predominantly localize to secondary lymphoid organs while effector Tregs are CD44^hi^ CD62L^lo^ CCR7^-^ and localize to both secondary lymphoid organs and non-lymphoid tissues (*30*). Here, we found altered Treg programming in both central and effector Foxp3-ΔE2 Treg populations, with the greatest impact on the effector Treg subset.

Interestingly, we identified an enrichment for T_H_17 signature genes in effector Foxp3-ΔE2 Tregs. Prior studies had hypothesized that Foxp3-ΔE2 Tregs would have increased expression of T_H_17 associated genes since Foxp3 Exon 2 is known to bind and suppress RORγt (*13*) but had not found evidence to support this hypothesis (*15, 19*). The enrichment of T_H_17 signature genes suggests impaired RORγt regulation by Foxp3-ΔE2 Tregs and may also reflect reduced IL-2/STAT5 signaling in Foxp3-ΔE2 Tregs which is known to suppress T_H_17 transcriptional programming (*31*). By separating central and effector Treg populations, we were able to identify the differential expression of T_H_17 signature genes in Foxp3-ΔE2 effector Tregs, highlighting the importance of parsing out these populations in analysis of Foxp3-ΔE2 Tregs.

The function of the FOXP3-ΔE2 isoform in humans remains unknown, despite approximately equivalent expression levels of the two FOXP3 isoforms in healthy individuals (*11*). FOXP3 Exon 2 spans amino acids 71-105; encodes binding sites for RORα, RORγt, and hnRNPF; and contains one of the two nuclear export signals found in FOXP3 (*12–14, 32, 33*). The precise molecular mechanisms underlying the differences in function between the two isoforms also remain unknown although they have largely overlapping DNA binding sites (*15*), suggesting that altered protein-protein complex formation by the Foxp3-ΔE2 isoforms may play a larger role in driving altered cellular programming and function. The signaling factors which regulate alternative splicing of FOXP3 remain unknown although inhibition of glycolysis favors FOXP3-ΔE2 expression (*34*). Clinical studies have uncovered a pathogenic role for the FOXP3-ΔE2 isoform in autoimmunity, with sole FOXP3-ΔE2 expression driving lethal IPEX syndrome and upregulation of FOXP3-ΔE2 in several autoimmune diseases (*15–18*).

Given the evident danger of FOXP3-ΔE2 expression, there is likely some evolutionary advantage to having this isoform. Our data suggest a possible benefit to alternative FOXP3 isoform usage in infection. During acute infection, increased FOXP3-ΔE2 usage could permit increased CD8^+^ T cell responses. Then, upon pathogen clearance, conversion to FOXP3-FL isoform usage could favor contraction of the response. This hypothesis is supported by the recent report that increased FOXP3-ΔE2 expression positively correlated with longer survival time in triple negative breast cancer, highlighting the potential benefit of FOXP3-ΔE2 expression in enhancing CD8^+^ T cell responses (*21*). Additionally, despite their suppressive defects, Foxp3-ΔE2 Treg numbers increase to a greater degree than Foxp3-FL Tregs in homozygous mice, likely due to their enhanced proliferative capacity. Therefore, FOXP3-ΔE2 expression could enhance Treg expansion during infection, leaving behind a larger pool of Tregs to then contract the immune response and contribute to wound healing. Future experiments should assess these possibilities to better understand FOXP3-ΔE2 Tregs in both health and disease.

The role of IL-2 signaling in CD8^+^ T cell responses and the regulation of this signaling by Tregs has proven controversial. Early work using OT-I CD25 KO CD8^+^ T cells concluded that IL-2 signaling through IL-2Rα was dispensable for early CD8^+^ T cell expansion in secondary lymphoid organs but was important for sustaining CD8^+^ T cell expansion in nonlymphoid tissues (*35*). However, recent work has demonstrated that IL-2 signaling during a secondary priming phase in the lymph node promotes CD8^+^ T cell expansion and effector differentiation (*36*). Plausibly, IL-2 signaling received in secondary lymphoid organs may support the expansion of CD8^+^ T cells that migrate into nonlymphoid tissues. This hypothesis is supported by our findings, where reduced co-localization of Foxp3-ΔE2 Tregs with CD8^+^ T cells and reduced IL-2 sink activity in the secondary lymphoid organ during priming drove an enhanced magnitude of the CD8^+^ T cell response in both lymph node and tissue.

Despite an enhanced magnitude of the CD8^+^ T cell response in Foxp3-ΔE2 mice during IAV infection, there was no evidence of enhanced viral clearance or worsened immunopathology. These data are likely explained by the lack of difference in the number of CD8^+^ short lived-effector cells that mediate viral clearance and cell killing between Foxp3-ΔE2/ΔE2 and Foxp3-FL/ΔE2 mice. Rather, it appears that the presence of Foxp3-ΔE2 Tregs skews the CD8^+^ T cell response to increased memory formation. Although we observed no difference in the function of these memory CD8^+^ T cells at an early memory timepoint in our heterosubtypic infection model, it is possible that they would have an enhanced protective function at later memory timepoints due to their enhanced long-term persistence. These data were initially unexpected as Tregs have been shown to selectively inhibit the generation of short-lived effector cells while preserving the induction of central memory cells in murine infection models with acute Treg depletion (*37, 38*) and IL-2 signaling in CD8^+^ T cells promotes effector differentiation (*39*). However, recent work using an IL-2 mutein demonstrated that enhanced IL-2 signaling during acute infection *in vivo* actually increases CD127 expression on CD8^+^ T cells (*40*). Since increased CD127 expression on CD8^+^ T cells is associated with enhanced CD8^+^ T cell memory formation and persistence (*41*), these data support a model where enhanced IL-2 signaling during the acute phase of infection may in fact drive increased CD8^+^ T cell memory formation. Alternatively, there may be an additional altered function in Foxp3-ΔE2 Tregs independent of IL-2 that leads to enhanced memory precursor formation in Foxp3-ΔE2 mice. We concluded that Tregs and/or IL-2 signaling may play a more complicated role than previously appreciated in CD8^+^ T cell effector and memory differentiation and that Foxp3-ΔE2 Tregs favor CD8^+^ T cell memory formation.

Spatial dynamics play a key role in the regulation of CD8^+^ T cell responses by Tregs. The initial phase of CD8^+^ T cell priming involves recruitment of naïve T cells to antigen presenting cells within the T cell zone of secondary lymphoid organs, and Tregs function as an IL-2 sink while dynamically traversing clusters of CD8^+^ T cells during priming (*36*). In this study, we found reduced localization of Foxp3-ΔE2 Tregs to the T cell zone of the lung-draining lymph node during T cell priming. This was driven by altered chemokine receptor expression on Foxp3-ΔE2 Tregs, including increased CCR7 and decreased CXCR3 expression. Increased CCR7 expression on Foxp3-ΔE2 Tregs might initially suggest increased localization to the T cell zone, which has high levels of the CCR7 ligands CCL19 and CCL21 under homeostatic conditions (*42*). However, during inflammatory responses, CCL21 is downregulated and the CXCR3 ligand, CXCL9, is upregulated (*25–29*). Concordantly, we found decreased CCL21 and increased CXCL9 expression in the lung-draining lymph node compared to the non-draining lymph node following influenza viral infection. Thus, Foxp3-ΔE2 Tregs have decreased presence in the T cell zone despite increased CCR7 expression. Additionally, Foxp3-ΔE2 Tregs were less poised for recruitment to the CXCL9-high T cell zone due to low CXCR3 expression. Together, these data suggest that altered chemokine receptor expression on Foxp3-ΔE2 Tregs led to reduced occupancy of the T cell zone, contributing spatially to impaired regulation of CD8^+^ T cells during priming.

A technical limitation of this study was that the rodent Foxp3-ΔE2 isoform is germline encoded while in humans it is produced through alternative splicing, so these mice do not allow us to assess the role that alternative splicing of Foxp3 may play in balancing Foxp3 isoforms and understanding how this alters regulation of conventional T cell responses. Additionally, there are many known mechanisms of conventional T cell regulation by Tregs (*32, 43, 44*). While we identified that defects in IL-2 regulation and impaired localization by Foxp3-ΔE2 Tregs contribute to aberrant CD8^+^ T cell regulation, the altered transcriptional programming of central and effector Foxp3-ΔE2 Tregs makes possible additional defective mechanisms of suppression by Foxp3-ΔE2 Tregs. Finally, in our Foxp3-DTR/ΔE2 mouse experiments, more Foxp3-DTR expressing Tregs are depleted in Foxp3-DTR/ΔE2 than Foxp3-DTR/FL mice since Foxp3-FL expressing Tregs outcompete Foxp3-ΔE2 Tregs. However, Foxp3-ΔE2 Tregs rapidly filled the Treg niche upon DT treatment, and there is no alternative model to acutely deplete WT Tregs to generate Foxp3-ΔE2 mice without prior background inflammation, making this model the best option for these experiments despite this potential caveat. Altogether, our data suggest that the Foxp3-ΔE2 isoform may act as a natural ‘dominant negative’ to precisely tune pathogen-specific immune responses, and future studies are needed to assess Foxp3 isoform usage in the context of infection.

Taken together, this study demonstrates that Foxp3 Exon 2 is required for optimal effector Treg programming, including expression of the high affinity IL-2Rα and tissue-homing chemokine receptors. Deficiency in expression of these molecules results in reduced IL-2 sequestration and localization to the T cell zone during priming, driving an enhanced magnitude of the CD8^+^ T cell response. These are the most comprehensive data to date describing central and effector Foxp3-ΔE2 Treg populations and the mechanisms by which Foxp3-ΔE2 Tregs promote CD8^+^ T cell responses, providing critical insights into altered immune regulation by Foxp3-ΔE2 Tregs which are broadly relevant in both human health and autoimmunity.

## MATERIALS AND METHODS

### Study design

The overall goal of this study was to identify phenotypic and functional differences in Tregs programmed by the FOXP3-ΔE2 isoform. We first studied central and effector Tregs in Foxp3-ΔE2 mice at steady state and during influenza infection. We next studied CD8^+^ T cell regulation in influenza infected Foxp3-ΔE2 mice then crossed our Foxp3-ΔE2 mice with Foxp3-DTR mice to study the cell-intrinsic nature of the defects in CD8^+^ T cell suppression by Foxp3-ΔE2 Tregs. We tested the function of *Il2ra* and *Ctla4* on Foxp3-ΔE2 Tregs *ex vivo* using IL-2/pSTAT5 signaling and a CD86-eGFP transendocytosis assay and *in vivo* using our influenza infection model. Finally, we studied co-localization of Foxp3-ΔE2 Tregs with conventional T cells in the lung-draining lymph node during influenza infection using confocal microscopy. The number of mice used and statistical tests performed are included in the figure legends, power calculations were performed following pilot experiments to determine appropriate sample sizes, and each experiment was performed 2-5 times.

### Mice

Foxp3-ΔE2 mice were generated as previously described(*15*). Strains C.B6-Tg(Foxp3-HBEGF/EGFP)23.2Spar/Mmjax (common name DEREG), B6.129(Cg)-*Foxp3^tm3(Hbegf/GFP)Ayr^*/J (common name Foxp3^DTR^), and C57BL/6-*Foxp3^tm1Flv^*/J (common name FOXP3-IRES-mRFP) were obtained from The Jackson Laboratory. All mice for this study were bred and maintained under specific pathogen-free conditions in an American Association for the Accreditation of Laboratory Animal Care (AAALAC)-accredited animal facility at the Benaroya Research Institute (BRI). All experiments were performed in accordance with protocols approved by the BRI Institutional Animal Care and Use Committee. Age, sex-matched, and littermate controls aged 8-12 weeks were used wherever possible. Some experiments required exclusive use of male or female mice because Foxp3 is located on the X chromosome and are explicitly labeled when applicable.

### Influenza virus

X31 influenza A virus (IAV) was generously provided by Dr. Andrew Oberst at the University of Washington. A/PR/8/34 IAV was purchased from ATCC (#VR-95). Mice were anesthetized with inhaled isoflurane then inoculated with virus by oropharyngeal inoculation at indicated doses. Mouse weight was measured once daily until recovered to baseline then measured once weekly.

### Tissue processing and cell isolation

In experiments where only secondary lymphoid organs were collected, mice were euthanized by CO_2_ inhalation followed by cervical dislocation. In experiments where lungs were collected, mice were anesthetized with a sub-lethal dose of i.p. injected tribromoethanol (TBE), retro-orbital injected with 2μg/mouse CD45-BUV395 to label vascular CD45^+^ cells, given 3-5 minutes for the antibody to circulate, and euthanized by thoracotomy. The left lung lobe was inflated with 10% neutral buffered formalin then stored in formalin at room temperature for histology, the right lung lobe was stored in supplemented RPMI on ice for flow cytometry, and the post-caval lobe was stored in RNAlater (Invitrogen #AM7021) at -80°C for RNA isolation. The mediastinal lymph node (medLN) and spleen were stored in RPMI on ice for flow cytometry. Tissues for flow cytometric analysis were processed to single cell suspensions as previously described, except the lung RBC incubation was performed for only 3 minutes.

### Flow cytometric analysis

For cytokine staining, cells were stimulated with phorbol 12-myristate 13-acetate (PMA) (10ng/mL) and ionomycin (500ng/mL) for 4 hours. Brefeldin A (Invitrogen #00-4506-51) was added after 1 hour of stimulation. Cells were stained with Zombie NIR Fixable Viability Kit for 10 minutes at 4°C (Biolegend #423106), Fc blocked (BioXCell #BE0307) for 5 minutes at room temperature, and surface stained with antibodies at 4°C for 30 minutes. In influenza infection experiments, cells were additionally surface stained at room temperature for 45 minutes with Class I MHC Tetramer H-2D^b^/Influenza A nucleoprotein (NP)_366-374_ (ASNENMETM) tetramer (National Institutes of Health Tetramer Core Facility, Emory University). Cells were either fixed and permeabilized with Cytofix/Cytoperm Fixation/Permeabilization kit (BD #554714) for cytokine staining or Foxp3/Transcription Factor Staining Buffer Set (eBioscience #00-5523-00) for transcription factors at 4°C for 30 minutes. Intracellular staining was performed at room temperature for 30 minutes. For pSTAT5 staining, cells were stained with viability dye then fixed in Phosflow Lyse/Fix Buffer (BD #558049) at 37°C for 10 minutes followed by 90% methanol for 30 minutes on ice. Cells were then permeabilized with Foxp3/Transcription Factor Staining Buffer Set permeabilization buffer and stained with antibodies for 45 minutes at room temperature. AccuCheck Counting Beads (Invitrogen #PCB100) were added to each sample to allow cell count enumeration. Data was acquired on the Cytek Aurora spectral flow cytometer and analyzed in FlowJo v10.10.0. Full antibody information is available in Supp Table 5.

### Histological analysis

Fixed lungs were imbedded in paraffin, sectioned, and stained with hematoxylin and eosin. Blind scoring of tissue pathology was performed by Dr. Jessica Snyder based on perivascular and periobronchiolar thickening, bronchiolar epithelium inflammation and hyperplasia, alveolar inflammation, interstitial inflammation, intrabronchiolar inflammation, and total extent of inflammation. Individual scores were summed to create a total lung pathology score.

### RNA isolation, cDNA synthesis, and qRT-PCR

Lung samples were weighed then transferred to a sterile 1.5mL tube, resuspended in 350uL buffer RLT (Qiagen #79216) with 1% beta-mercaptoethanol, disrupted using the BioMasher Standard attachment (TaKaRa #9791A) in the PowerMasher II (Nippi #891300), and homogenized using a 1mL syringe fitted with a 18G needle. Samples were centrifuged at 3000rpm for 3 minutes and the supernatant was processed for RNA isolation using the RNeasy Mini Kit (Qiagen #74104). RNA was reverse transcribed with a High-Capacity cDNA Reverse Transcription Kit (AppliedBiosystems #4368814) following manufacturer’s protocol. qRT-PCR was prepared using TB Green Premix Ex Taq II (TaKaRa #RR82WR) following manufacturer’s protocol and data was acquired on an Applied Biosystem 7500 Fast Real-Time PCR System.

### Cell culture

Primary mouse cells and CD86-eGFP transduced 3T3 fibroblasts (*22*) were cultured in RPMI-1640 (Cytiva #SH30255.01) containing 25mM HEPES and L-Glutamine then supplemented with 10% FBS (Sigma-Aldrich #F4135), 1mM sodium pyruvate (Cytivia #SH40003.01), 1X MEM non-essential amino acids (Gibco #11140-050), and 1X beta-mercaptoethanol (Gibco #21985-023). For IL-2 stimulation experiments, bulk populations of cells were isolated from lungs as described above or CD4^+^ T cells were enriched from naïve mouse spleens using an EasySep Mouse CD4^+^ T cell Isolation kit following manufacturer’s protocol (StemCell #19852). IL-2 recombinant protein (Gibco #PHC0021) was added to cells at indicated concentrations and cells were incubated at 37°C for 30 minutes prior to staining for analysis by flow cytometry. For the transendocytosis assay, primary mouse CD4^+^ T cells were cultured with CD86-eGFP 3T3 cells at a 1:1 ratio for 3 hours at 37°C. All wells were cultured with 25nM Bafilomycin A1 (Sigma-Aldrich #88899-55-2) and, where indicated, cells were cultured with 100µg/mL α-CTLA-4 (Bio X Cell #BE0032) and/or ImmunoCult Mouse T cell Activator (StemCell #100-1572). Non-adherent cells were harvested and analyzed for CD86 internalization by flow cytometry.

### Immunofluorescence

Dual reporter Foxp3-FL-mRFP/Foxp3-ΔE2 DEREG^+^ mice were infected with IAV and the mediastinal and inguinal lymph nodes were dissected at 3DPI. Isolated LN tissues were fixed using BD Cytofix (BD Biosciences #554655), diluted 1:3 in PBS for 20–24 h at 4°C, and dehydrated with 30% sucrose solution for 24–48 h at 4°C. LNs were then embedded in an OCT compound (Tissue-Tek #4583) and stored at −20°C. LNs were sectioned on a Thermo Fisher Scientific Micron HM550 cryostat into 20-µm sections and stained as previously described (*45*). A Leica SP8 tiling confocal microscope equipped with a 40× 1.3 NA oil objective was used for image acquisition. All raw imaging data was processed and analyzed using Imaris (Bitplane).

### Bulk RNA-seq

For each cell subset, 200 cells were sorted directly into reaction buffer using the SMART-seq v4 Ultra Low Input RNA Kit for Sequencing (Takara). Reverse transcription and PCR amplification were performed to generate full-length amplified cDNA and sequencing libraries were constructed using the NexteraXT DNA sample preparation kit (Illumina). Paired end reads at a run length of 59 nucleotides were acquired on an Illumina NextSeq 2000 with a target depth of 5 million reads per sample. Base calls were processed to FASTQs on BaseSpace (Illumina), and, using trimmomatic (v0.39), a base call quality-trimming step was applied to remove low-confidence base calls from the ends of read. The FASTQs were aligned to the mouse reference genome (GRCm38.91) using STAR (v.2.7.11a). After identification (Picard v3.1.0) and removal (samtools v1.16) of duplicates, gene counts were generated using htseq-count (v2.0.2). QC and metrics analysis was performed using fastQC (v0.12.1) and Picard (v3.1.0). Differential gene expression was calculated using DESeq2 in R (v4.4.2). PCA plots were generated using pcaExplorer (v3.0.0), and heat maps and volcano plots were created in R using custom code. Gene set enrichment analysis (GSEA) was performed from total raw gene count lists using the GSEA 4.3.2 application.

### Statistical analysis

Sample sizes were determined based on a two-tailed power calculation using an error rate of 0.05, power of 80%, and estimated effect size and variability from preliminary data. Statistical analysis was performed in GraphPad Prism v10.4.1 and presented as mean ± standard deviation. Unpaired or paired t tests, multiple unpaired or paired t tests with Holm-Sidak method of correction for multiple comparisons, and one-way or two-way analysis of variance (ANOVA) were used for hypothesis testing. A generalized additive mixed model was used to compare weight loss curves. P < 0.05 was considered statistically significant. ns (not significant): P > 0.05, * P < 0.05, ** P < 0.01, *** P < 0.001, and **** P < 0.0001.

## Supplementary Materials

Table. S1. Differentially expressed genes between central Foxp3-FL vs ΔE2 Tregs.

Table. S2. Differentially expressed genes between effector Foxp3-FL vs ΔE2 Tregs.

Table. S3. Differential gene expression between total Foxp3-ΔE2 and Foxp3-FL Tregs.

Table. S4. Differential gene expression between lung parenchymal Foxp3-ΔE2 and FL Tregs at 7DPI with IAV.

Table. S5. Antibody list.

## Acknowledgments

We would like to thank Dr. Jessica Snyder for her technical expertise in scoring mouse lung pathology, the National Institutes of Health Tetramer Core Facility (NIH Contract 75N93020D00005 and RRID:SCR_026557) for providing the influenza A NP 366-374 tetramer, and the Benaroya Research Institute Cell and Tissue Analysis, Genomics, and Animal Core Facilities.

## Author contributions

Conceptualization: KNW, BZ, PPD, MYG, DC, SFZ

Experimentation: KNW, ZHB, EAS, FNB, GZ, ML, JRA, PPD

Data analysis: KNW, ZHB, EAS, GZ

Consulting statistician: AK

Funding acquisition: KNW, SFZ

Writing: KNW, DC, SFZ

## Data and materials availability

RNA-seq datasets are deposited to GEO and are available under accession number GSE324381. All data supporting the findings in this study are available upon request to the corresponding author.

---

**Fig. S1.**
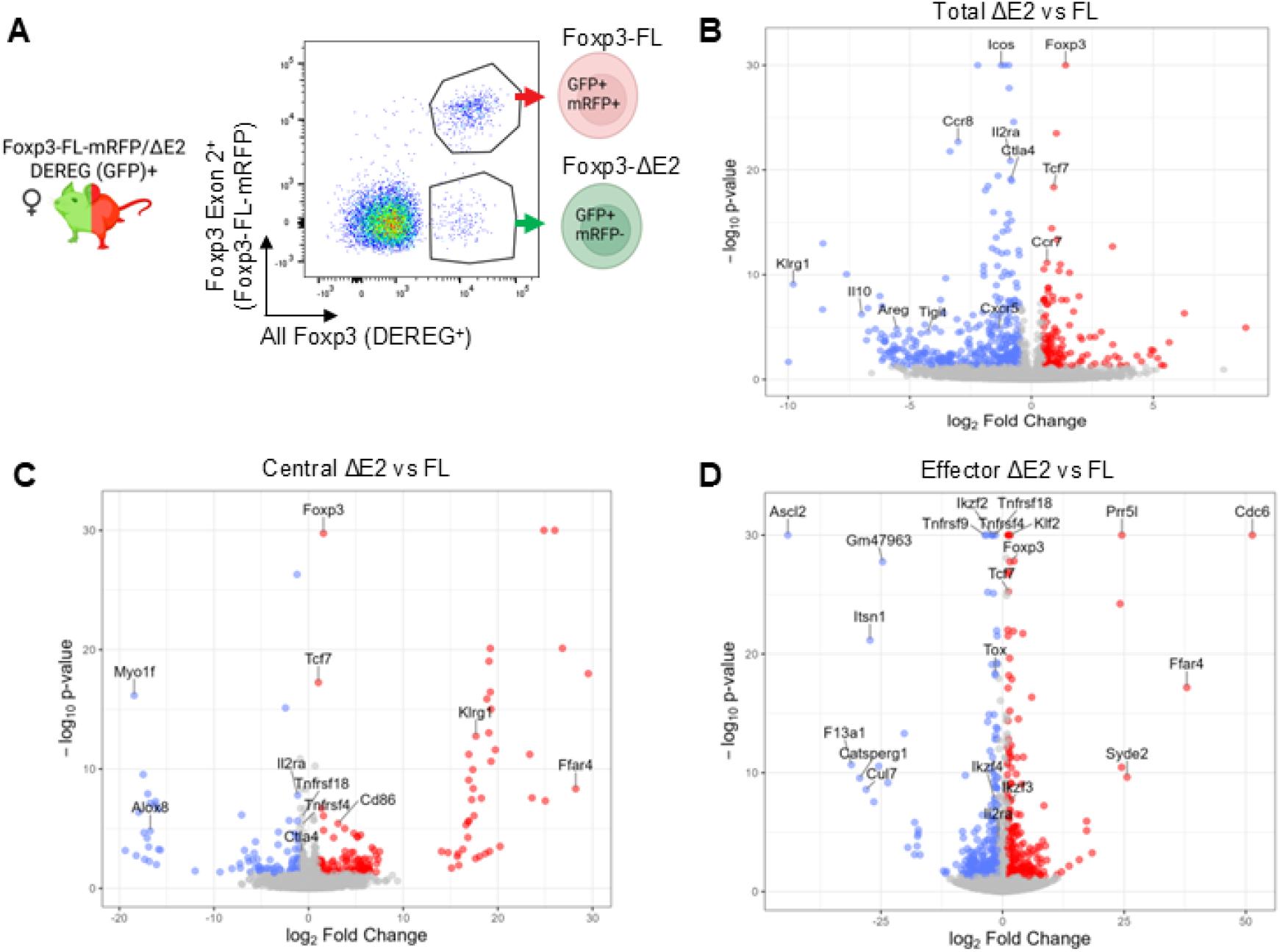
Differential gene expression in total, central, and effector Foxp3-ΔE2 Tregs. **(A)** A representative flow plot showing total Foxp3-FL and Foxp3-ΔE2 Tregs which were sorted from the spleens of dual reporter Foxp3-FL-mRFP/ΔE2 DEREG^+^ mice then analyzed by bulk RNA-seq. **(B)** Volcano plot showing differential gene expression between total Foxp3-ΔE2 vs Foxp3-FL Tregs sorted from dual reporter Foxp3-FL-mRFP/ΔE2 DEREG^+^ mouse spleens where red indicates upregulation and blue indicates downregulation in Foxp3-ΔE2 Tregs. **(C-D)** Volcano plots showing differential gene expression between **(C)** central or **(D)** effector Foxp3-ΔE2 vs FL Tregs sorted from dual reporter Foxp3-FL-mRFP/ΔE2 DEREG^+^ mouse spleens where red indicates upregulation and blue indicates downregulation in Foxp3-ΔE2 Tregs. *n*=5-6 mice.

**Fig. S2.**
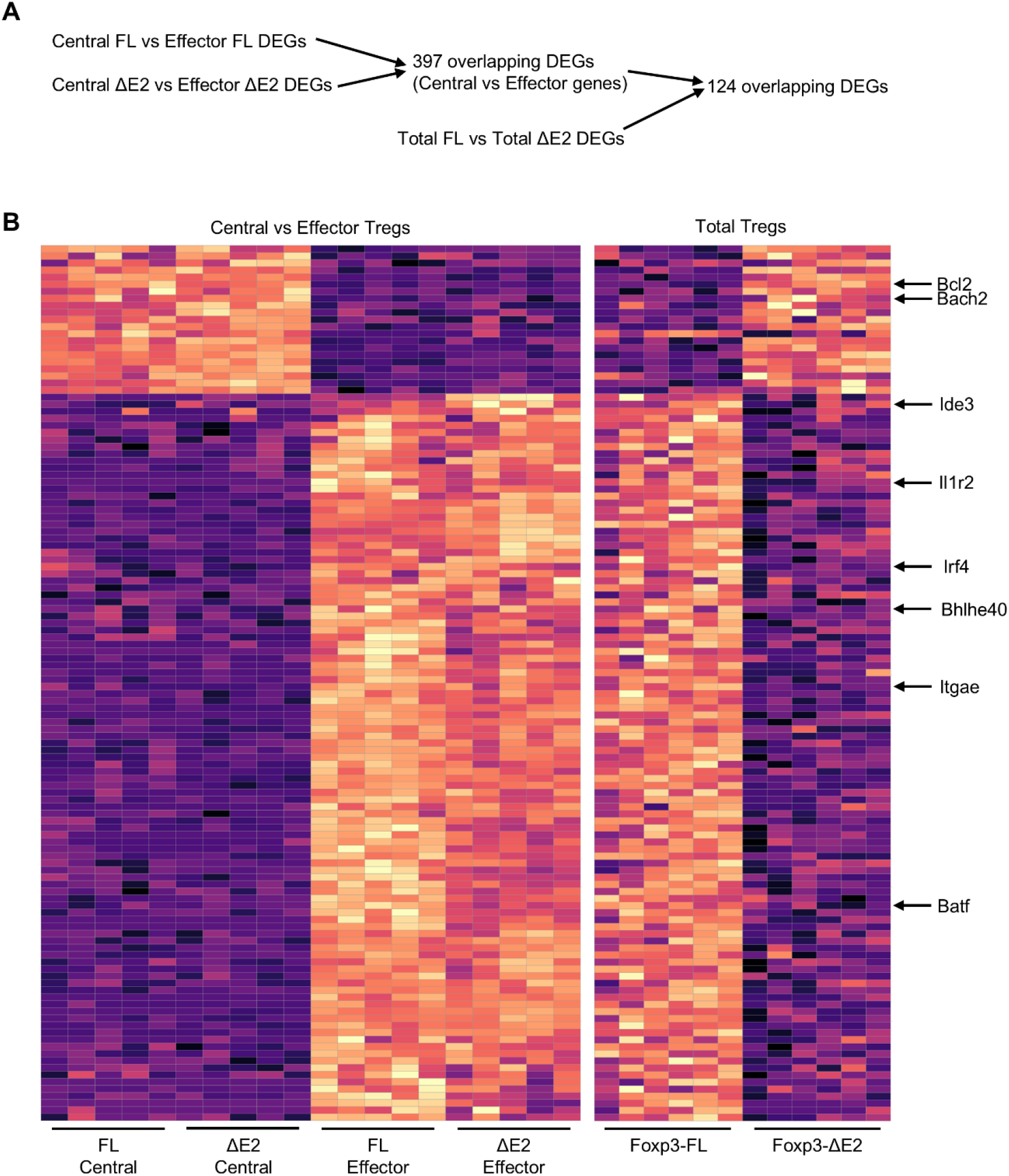
Foxp3-ΔE2 Treg DEGs driven by central Treg programming. **(A)** Diagram of DEG filtering. First, DEGs were identified for the separate comparisons of central Foxp3-FL vs effector Foxp3-FL Tregs and central Foxp3-ΔE2 vs effector Foxp3-ΔE2 Tregs then overlapped. These overlapping genes are the genes which are differentially expressed between central and effector Treg populations regardless of genotype. Next, total Foxp3-FL vs Foxp3-ΔE2 Treg DEGs were identified then overlapped with the central vs effector DEG list. The overall goal of this analysis was to identify which DEGs between total Foxp3-FL and Foxp3-ΔE2 Tregs are driven by central Treg programming, not isoform specific programming. **(B)** Z-scored heatmaps of the total Foxp3-FL and Foxp3-ΔE2 Treg DEGs which are driven by central Treg programming in Foxp3-ΔE2 Tregs shown for both total Foxp3-FL and Foxp3-ΔE2 Tregs and separately sorted central and effector Foxp3-FL and Foxp3-ΔE2 Tregs. *n*=5-6 mice per group.

**Fig. S3.**
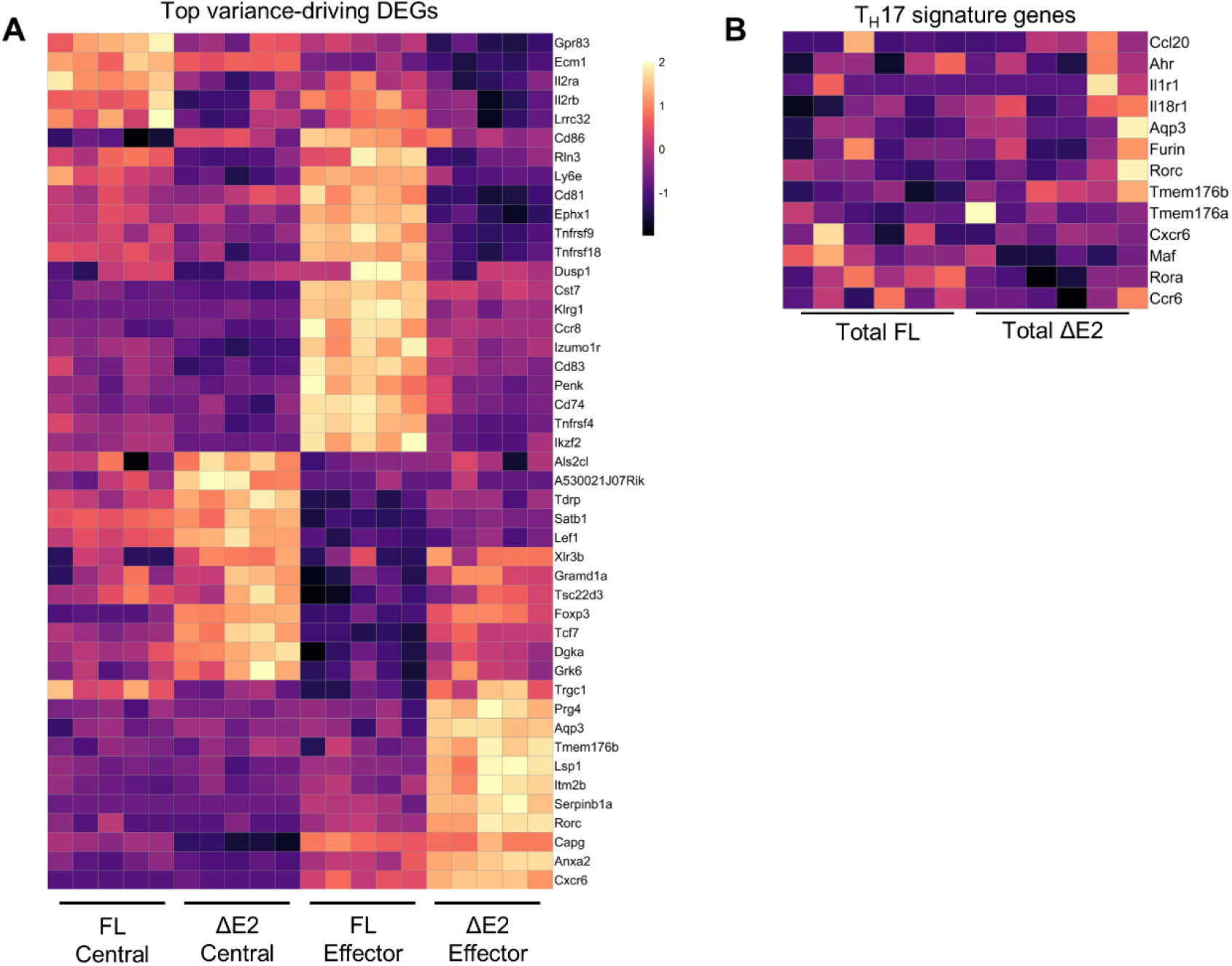
Altered cellular programming in central and effector Foxp3-ΔE2 Tregs. **(A)** A Z-scored heatmap of the top variance-driving differentially expressed genes between the four indicated Treg populations. **(B)** A Z-scored heatmap of T_H_17 signature genes in total Foxp3-FL and Foxp3-ΔE2 Tregs. *n*=5-6 mice.

**Fig. S4.**
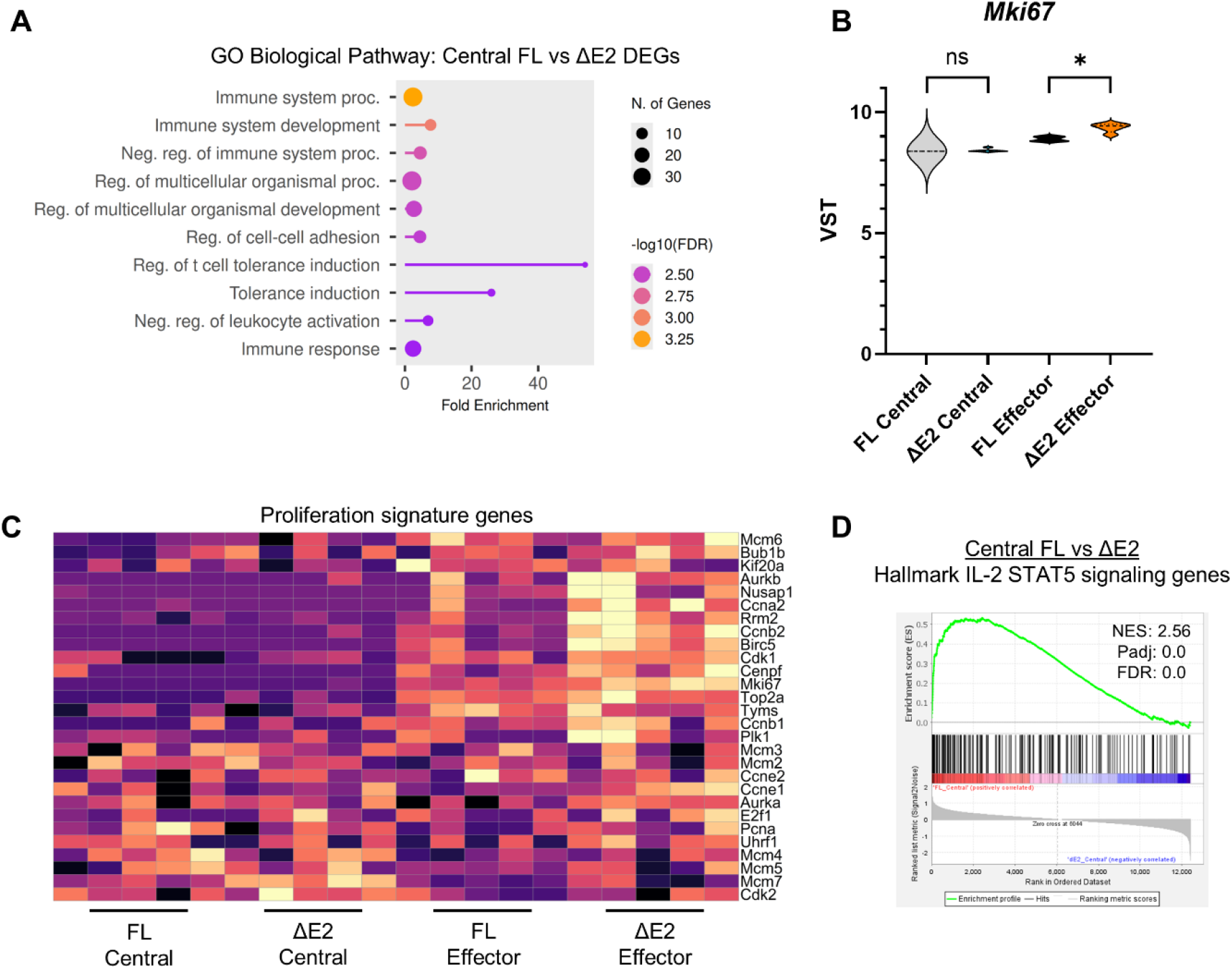
Pathway-level analyses of central and effector Foxp3-ΔE2 Tregs. **(A)** A GO Biological Pathway analysis of central Foxp3-FL vs Foxp3-ΔE2 Treg DEGs. **(B)** Violin plot showing VST *Mki67* expression for central and effector Foxp3-FL and Foxp3-ΔE2 Tregs. **(C)** A Z-scored heatmap of the proliferation signature genes in central and effector Foxp3-FL and Foxp3-ΔE2 Tregs. **(D)** GSEA of central Foxp3-FL vs Foxp3-ΔE2 Tregs for hallmark IL-2/STAT5 signaling genes with NES, padj, and FDR shown. *n*=5 mice. \**p*<0.05 by Wald Test performed within DESeq2 with adjusted p-value reported (B).

**Fig. S5.**
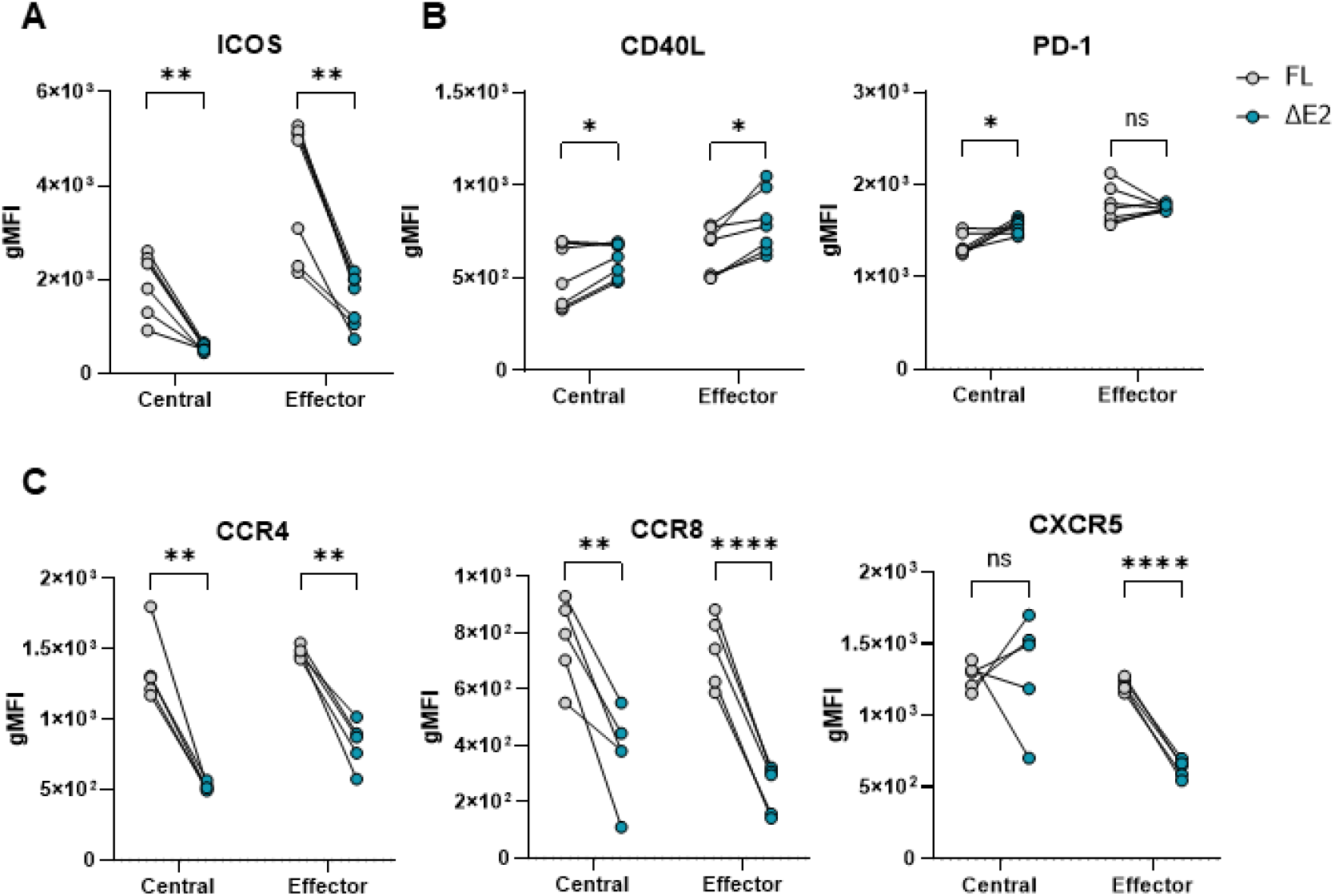
Altered expression of key functional molecules and chemokine receptors on central and effector Foxp3-ΔE2 Tregs. **(A-C)** Cell surface gMFI expression levels of (A) ICOS, (B) CD40L and PD-1, and (C) CCR4, CCR8, and CXCR5 on central and effector Foxp3-FL and Foxp3-ΔE2 Tregs from the spleens of steady-state Foxp3-FL/ΔE2 mice aged 8-12 weeks. *n*=5 mice. Pairing is shown within each mouse. \**p*<0.05, \*\**p*<0.01, \*\*\**p*<0.001, \*\*\*\**p*<0.0001 by two-tailed paired t test with Holm-Sidak method of correction for multiple comparisons (A-C).

**Fig. S6.**
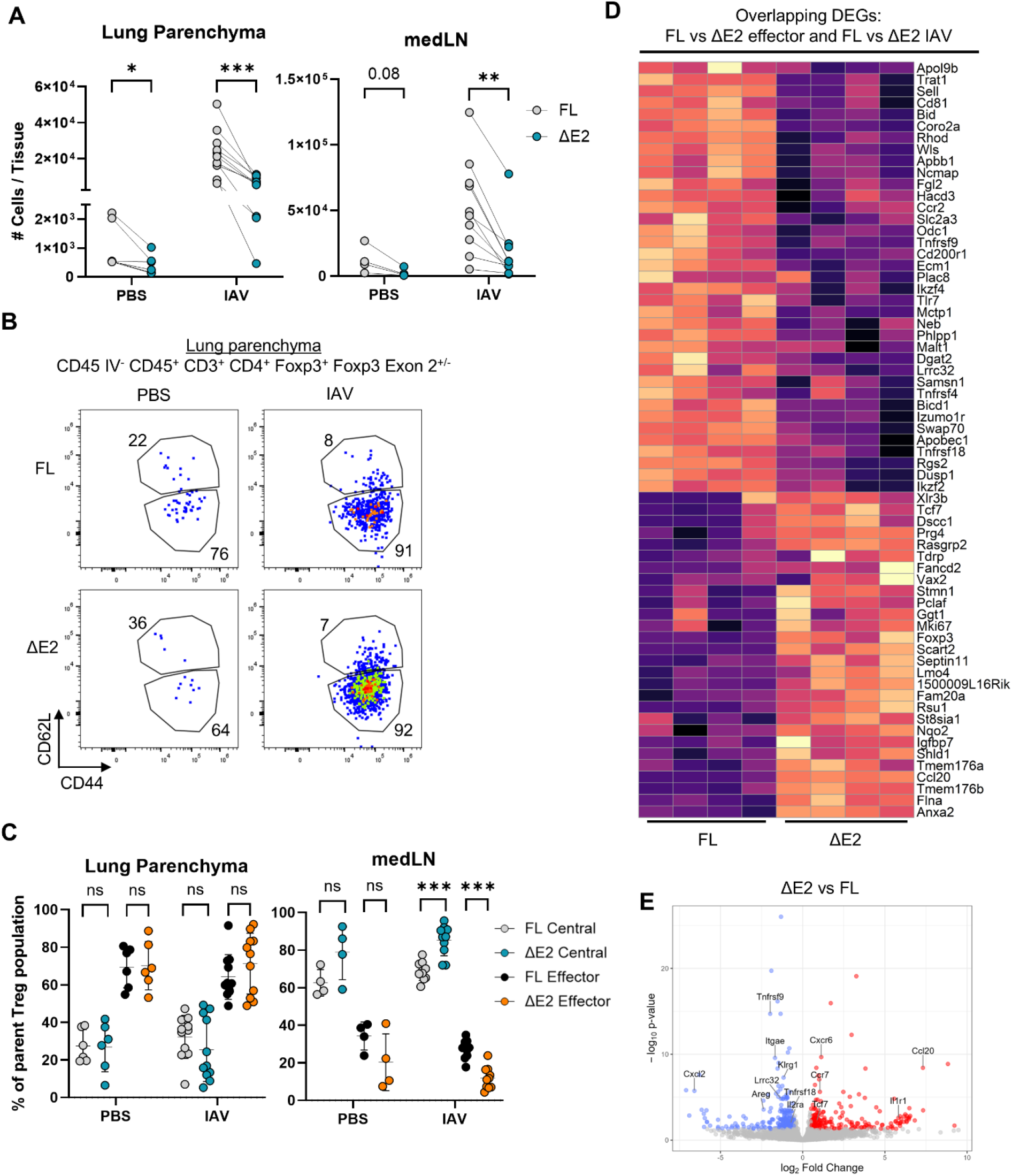
Central and effector Foxp3-ΔE2 Treg response to influenza viral infection. **(A-C)** Foxp3-FL/ΔE2 mice were PBS mock infected or infected with X31 IAV then analyzed by flow cytometry at 7DPI. **(A)** The numbers of Foxp3-FL and Foxp3-ΔE2 Tregs in the lung parenchyma and medLN. **(B)** Representative flow plots of central and effector Foxp3-FL and Foxp3-ΔE2 Tregs in the lung parenchyma. **(C)** The percentage of central and effector cells within total Foxp3-FL or Foxp3-ΔE2 Tregs. **(D-E)** Total Foxp3-FL and Foxp3-ΔE2 Tregs were sorted from the parenchyma of X31 IAV infected dual reporter Foxp3-FL-mRFP/ΔE2 DEREG^+^ mouse lung parenchyma at 7DPI then analyzed by bulk RNA-seq. **(D)** A Z-scored heatmap of DEGs which are differentially expressed between Foxp3-FL and Foxp3-ΔE2 effector Tregs at both homeostasis and during IAV infection. **(E)** A volcano plot of differentially expressed genes between Foxp3-ΔE2 vs Foxp3-FL Tregs where red indicates upregulation and blue indicates downregulation in Foxp3-ΔE2 Tregs. *n*=4 mice. \**p*<0.05, \*\**p*<0.01, \*\*\**p*<0.001 by multiple two-tailed paired t tests with Holm-Sidak method of correction for multiple comparisons (A & C).

**Fig. S7.**
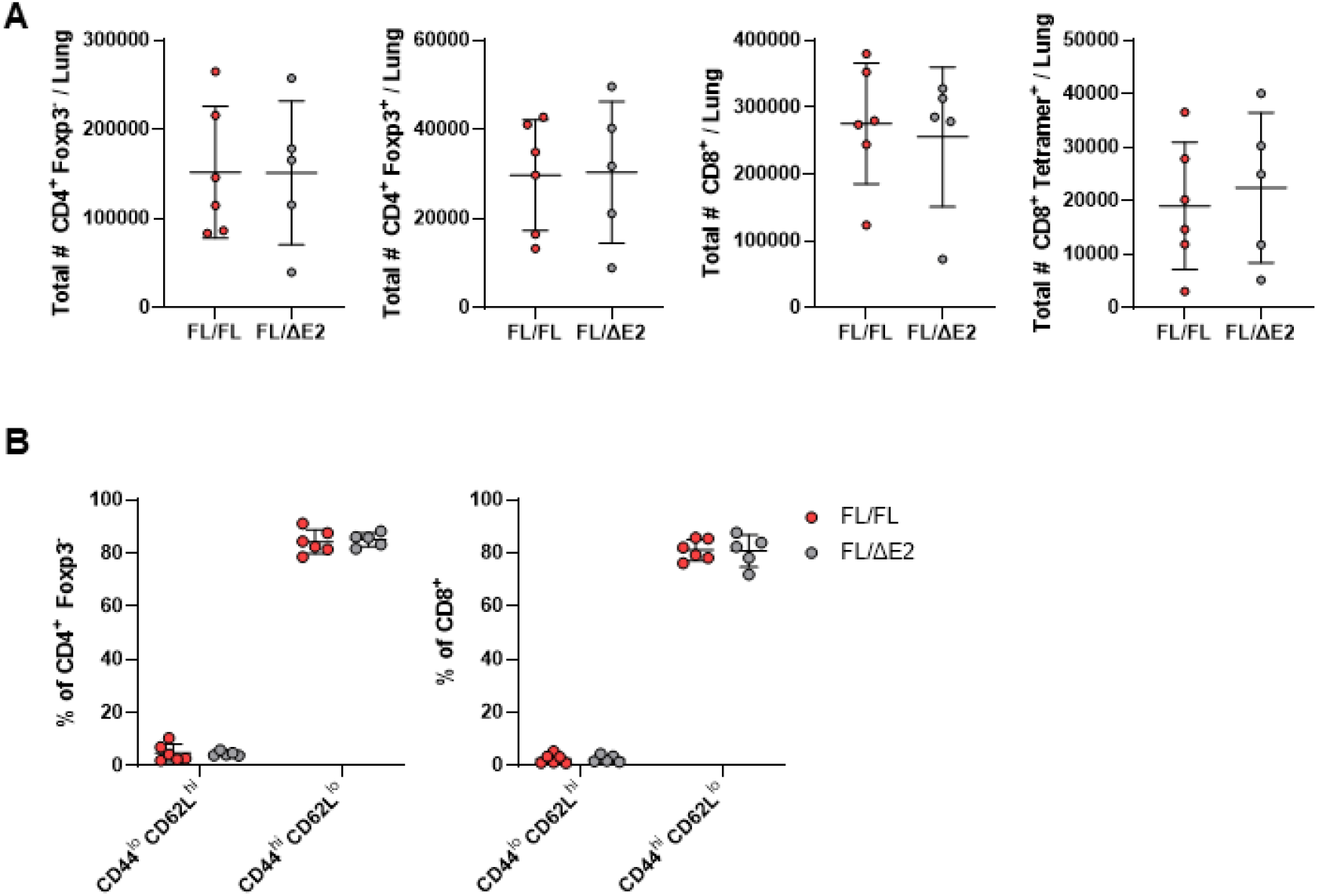
No difference in immune activation between IAV infected Foxp3-FL/ΔE2 and FL/FL mice. Foxp3-FL/ΔE2 and Foxp3-FL/FL (WT) mice were infected with X31 IAV and lung parenchyma localized immune cells were analyzed by flow cytometry at 7DPI. **(A)** The number of CD4^+^ Foxp3^-^, CD4^+^ Foxp3^+^, CD8^+^, and CD8^+^ NP-Tetramer^+^ cells. **(B)** The percentage of CD44^hi^ CD62L^lo^ and CD62L^hi^ CD44^lo^ cells amongst CD4^+^ Foxp3^-^ and CD8^+^ T cells. *n*=5-6 mice per group. Bars show mean values; error bars show SD.

**Fig. S8.**
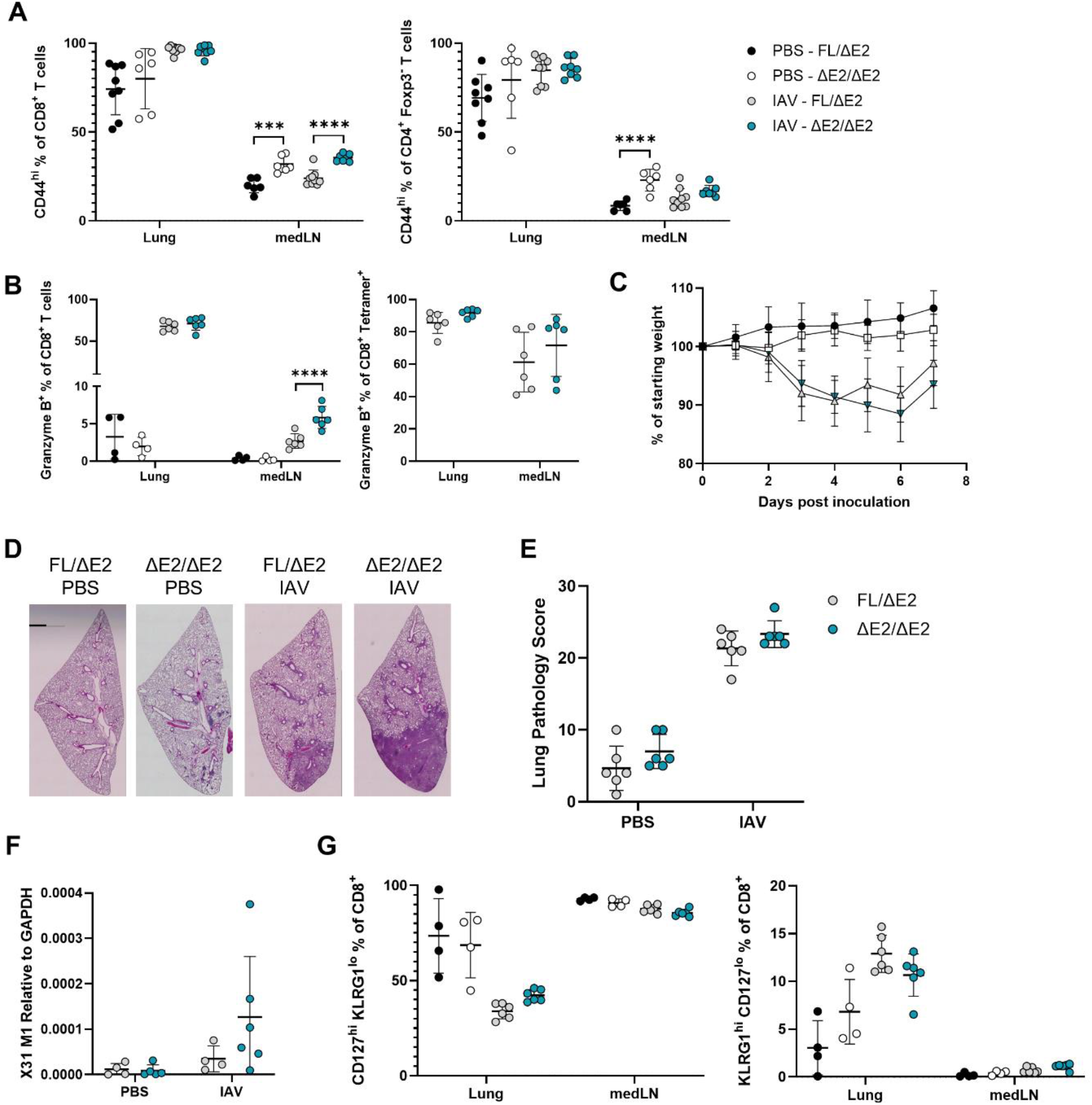
Increased CD8^+^ T cell response to acute IAV infection in Foxp3-ΔE2/ΔE2 mice. Foxp3-ΔE2/ΔE2 or littermate control Foxp3-FL/ΔE2 mice were PBS mock infected or infected with X31 IAV then analyzed during acute infection at 7DPI. **(A)** The percentage of CD8^+^ and CD4^+^ Foxp3^-^T cells which are CD44^hi^ in the lung parenchyma and medLN. **(B)** The percentage of Granzyme B^+^ cells amongst total CD8^+^ and CD8^+^ NP-Tetramer^+^ T cells. **(C)** Weight loss as a percentage of starting weight within each mouse. **(D)** Representative images of lung H&E staining. **(E)** Quantification of lung pathology from H&E staining. **(F)** qRT-PCR quantification of IAV using X31 M1 transcripts normalized to GAPDH. **(G)** The percentage of CD127^hi^ KLRG1^lo^ or KLRG1^hi^ CD127^lo^ cells amongst total CD8^+^ T cells. *n*=6-9 mice per group. Bars show mean values; error bars show SD. \*\*\**p*<0.001, \*\*\*\**p*<0.0001 by 2-way ANOVA followed by post-hoc t tests (A-B).

**Fig. S9.**
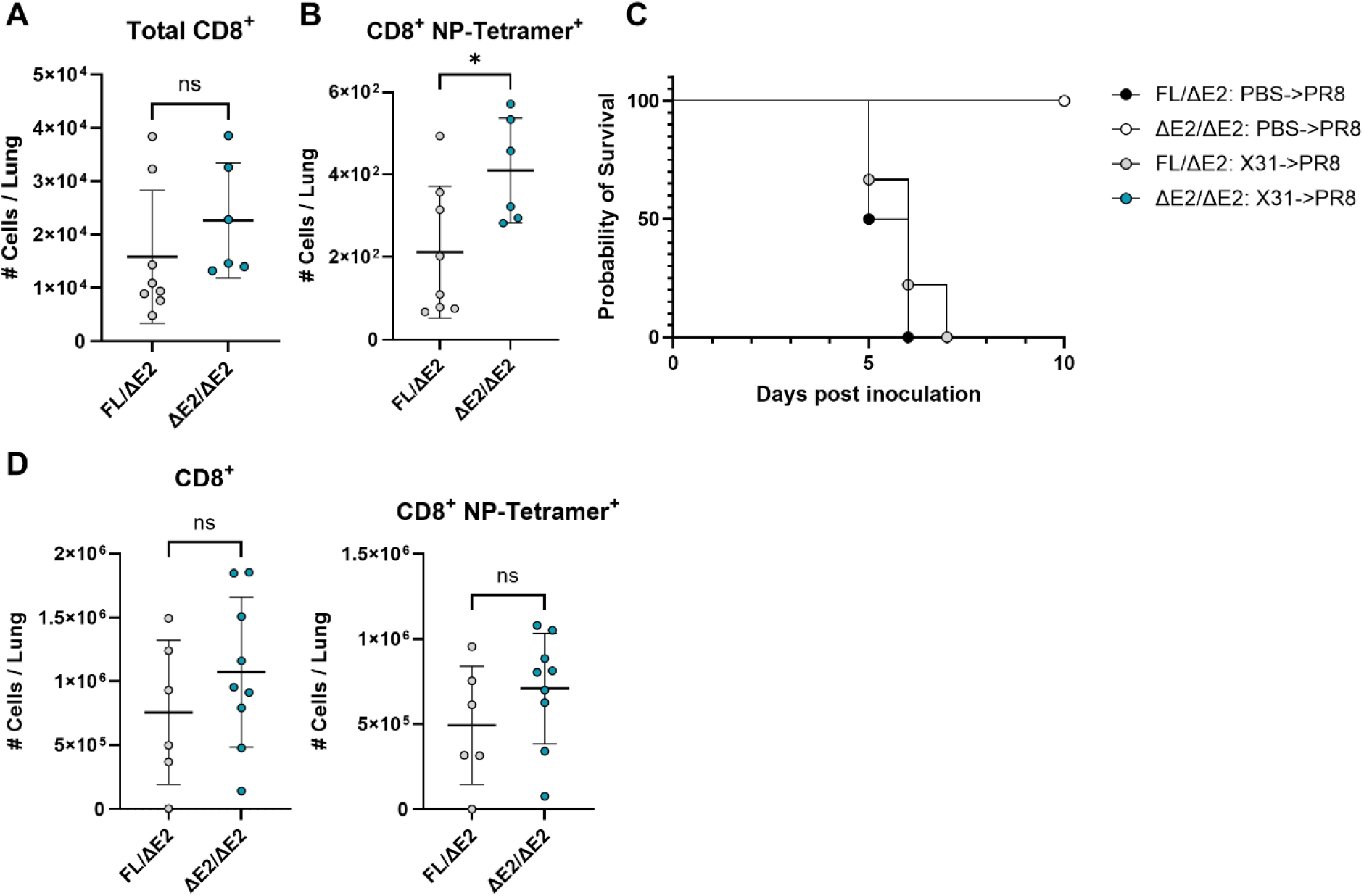
Increased CD8^+^ T cell memory formation following IAV infection in Foxp3-ΔE2/ΔE2 mice. **(A)** Foxp3-ΔE2/ΔE2 or littermate control Foxp3-FL/ΔE2 mice were infected with X31 IAV and analyzed during early memory at 30DPI. The number of total CD8^+^ T cells in the lung parenchyma is shown. **(B)** Foxp3-ΔE2/ΔE2 or littermate control Foxp3-FL/ΔE2 mice were infected with X31 IAV and analyzed during late memory at 90DPI. The number of CD8^+^ NP-Tetramer^+^ T cells is shown. **(C-D)** Foxp3-ΔE2/ΔE2 or littermate control Foxp3-FL/ΔE2 mice were either PBS mock infected or X31 IAV infected, recovered for 30 days to allow CD8^+^ T cell memory formation, then infected with a dose of PR8 IAV that is lethal in previously uninfected mice. **(C)** The probability of survival by day following PR8 IAV infection. **(D)** The number of total CD8^+^ and CD8^+^ NP-Tetramer^+^ T cells in the parenchyma of the lungs at 10DPI with PR8 IAV in surviving mice (FL/ΔE2 or ΔE2/ΔE2 mice which were X31→PR8 infected). *n*=6-9 mice per group. Bars shown mean; error bars show SD. \**p*<0.05 by unpaired two-tailed t tests.

**Fig. S10.**
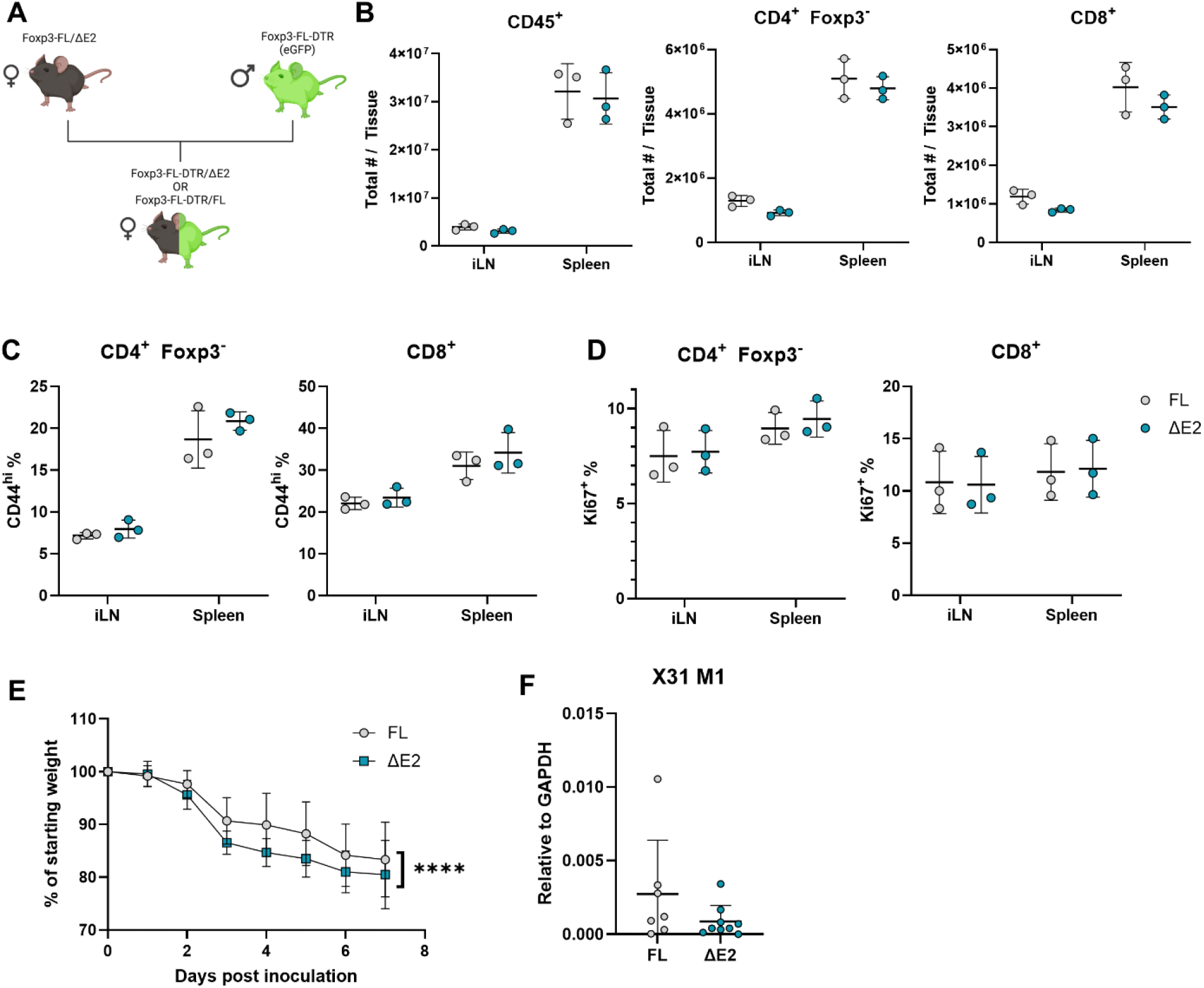
Foxp3-DTR x ΔE2 mice are a novel model to study Foxp3-ΔE2 Tregs without underlying systemic inflammation. **(A)** Schema of Foxp3-DTR x Foxp3-ΔE2 mouse breeding. **(B-D)** Female Foxp3-DTR/Foxp3-FL (FL) and Foxp3-DTR/Foxp3-ΔE2 (ΔE2) mice aged 8-12 weeks were analyzed for immune activation in the inguinal LN (iLN) and spleen by flow cytometry. **(B)** Total CD45^+^, CD4^+^ Foxp3^-^, and CD8^+^ T cell counts. **(C)** The percentage of CD44^hi^ cells amongst CD4^+^ Foxp3^-^ and CD8^+^ T cells. **(D)** The percentage of Ki67^+^ cells amongst CD4^+^ Foxp3^-^ and CD8^+^ T cells. **(E)** Foxp3-DTR x Foxp3-ΔE2 or littermate control Foxp3-DTR x Foxp3-FL mice were DT treated 2 days prior to infected, infected with X31 IAV, then DT treated every other day until analysis at 7DPI. Weight loss as a percentage of starting weight within each mouse is shown. **(F)** X31 M1 mRNA normalized to GAPDH by qRT-PCR from the post caval lung lobe at 7DPI with X31 IAV in DT treated Foxp3-DTR x ΔE2 mice. *n*=3-9 mice per group. Bars show mean; error bars show SD. \*\*\*\**p*<0.0001 by generalized additive mixed model test (E).

**Fig. S11.**
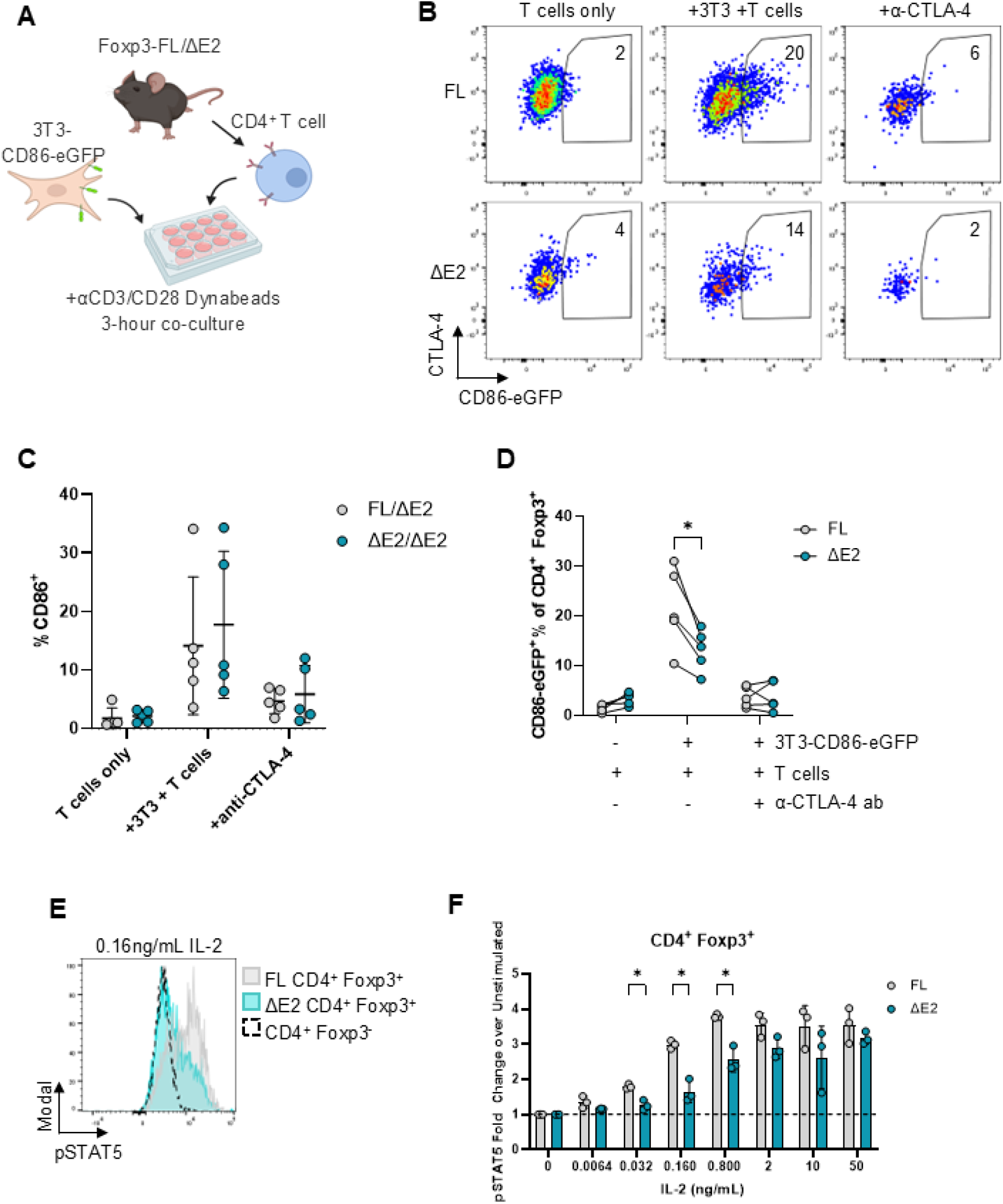
IL-2 and CD86 regulation by Foxp3-ΔE2 Tregs *ex vivo*. **(A-D)** CD4^+^ T cells from Foxp3-ΔE2/ΔE2 or FL/ΔE2 mice were MACS-enriched then co-cultured with CD86-eGFP expressing 3T3 cells to measure Treg transendocytosis of CD86 by flow cytometry. **(A)** Experiment schematic. **(B)** Representative flow plots show CD86-eGFP and CTLA-4 expression in T cells cultured alone, T cells co-cultured with 3T3-CD86-eGFP cells, and T cells co-cultured with 3T3-CD86-eGFP cells in the presence of an α-CTLA-4 blocking antibody. **(C)** Percentages of total Tregs from Foxp3-ΔE2/ΔE2 or FL/ΔE2 females which are CD86-eGFP^+^ following co-culture with 3T3-CD86-eGFP cells. **(D)** Percentages of Foxp3-FL or Foxp3-ΔE2 Tregs from Foxp3-FL/ΔE2 heterozygous females which are CD86-eGFP^+^ following co-culture with 3T3-CD86-eGFP cells. **(E-F)** CD4^+^ T cells were isolated from hemizygous Foxp3-FL or Foxp3-ΔE2 male mice, *ex vivo* stimulated with indicated concentrations of IL-2 for 30 minutes, and analyzed for pSTAT5 levels by flow cytometry. **(E)** A representative histogram of pSTAT5 induction in Foxp3-FL (gray), Foxp3-ΔE2 (cyan), or CD4^+^ Foxp3^-^T cells (black dashed line) upon stimulation with 0.16ng/mL IL-2. **(F)** Levels of pSTAT5 normalized to unstimulated cells within mouse at indicated concentrations of IL-2. *n*=3-5 mice per group. Bars show mean values; error bars show SD. \**p*<0.05 by multiple two-tailed unpaired t tests with Holm-Sidak method of correction for multiple comparisons (B) and two-tailed paired t tests with Holm-Sidak method of correction for multiple comparisons (F).

**Fig. S12.**
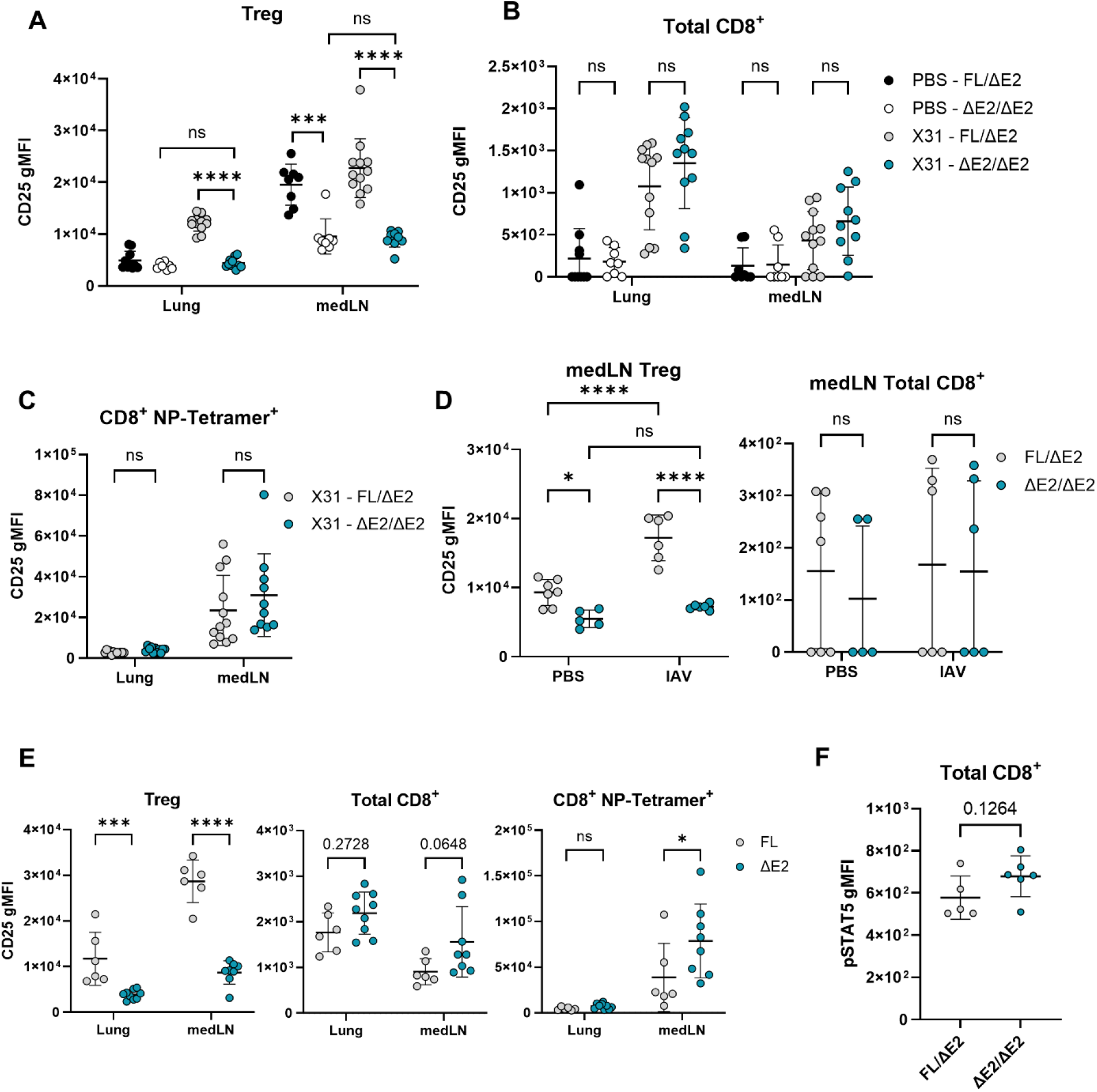
Deficiency in CD25 expression by Foxp3-ΔE2 Tregs during IAV infection. **(A-C)** Foxp3-ΔE2/ΔE2 and littermate control FL/ΔE2 mice were PBS mock infected or infected with X31 IAV and the lungs and medLN were analyzed at 7DPI by flow cytometry. CD25 gMFI on CD4^+^ Foxp3^+^ Tregs **(A)**, total CD8^+^ T cells **(B)**, and CD8^+^ NP-Tetramer^+^ cells from IAV infected mice only **(C)** is shown. **(D)** Foxp3-ΔE2/ΔE2 and Foxp3-FL/ΔE2 mice were X31 IAV infected and the medLN were analyzed at 3DPI by flow cytometry for CD25 gMFI expression on CD4^+^ Foxp3^+^ Tregs and total CD8^+^ T cells. **(E)** Foxp3-DTR x Foxp3-ΔE2 and Foxp3-DTR x Foxp3-FL mice were DT treated 2 days prior to infection to deplete Foxp3-DTR expressing Tregs, X31 IAV infected at day 0, then maintained with DT treatment every other day until analysis at 7DPI. CD25 gMFI expression on CD4^+^ Foxp3^+^ Tregs, total CD8^+^ T cells, and CD8^+^ NP-Tetramer^+^ cells from the lung parenchyma and medLN is shown. **(F)** Foxp3-FL/ΔE2 and Foxp3-ΔE2/ΔE2 mice were X31 IAV infected and CD8^+^ T cells from the medLN were analyzed for pSTAT5 gMFI expression. n=5-12 mice per group. Bars show mean values; error bars show SD. \**p*<0.05, \*\*\**p*<0.001, \*\*\*\**p*<0.0001 by two-way ANOVA followed by multiple t tests (A-B) or two-tailed unpaired t tests followed by Holm-Sidak method of correction for multiple comparisons where applicable (C-F).

**Fig. S13.**
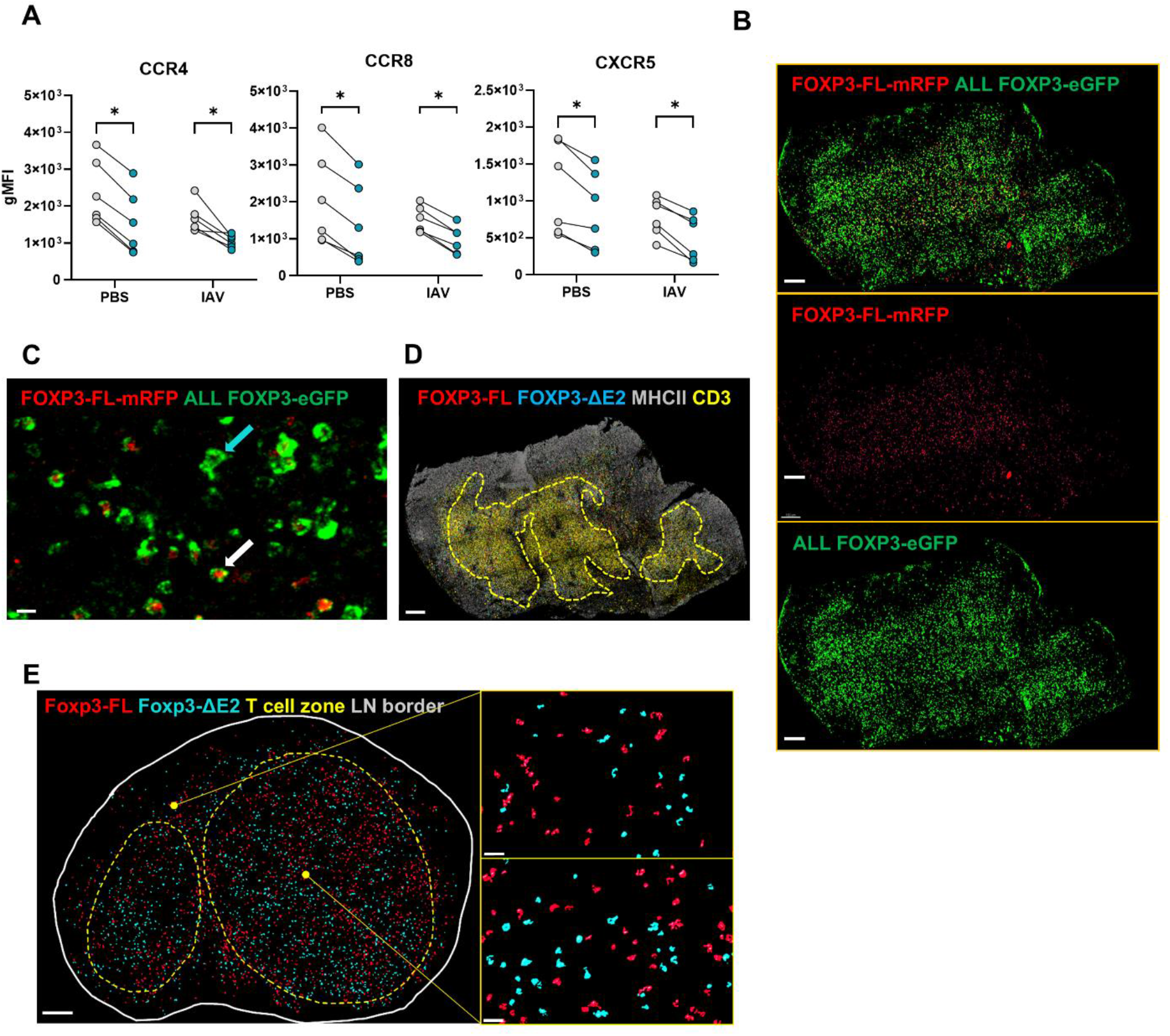
Altered Foxp3-ΔE2 Treg localization in secondary lymphoid organs. **(A)** Foxp3-FL/ΔE2 mice were X31 IAV infected and chemokine receptor gMFI expression was measured on CD4^+^ Foxp3^+^ Tregs in the medLN at 3DPI. **(B-E)** Foxp3-FL-mRFP/ΔE2 DEREG^+^ dual reporter mice were infected with IAV and the draining (mediastinal) and non-draining (inguinal) LNs were analyzed for Treg localization by confocal microscopy. **(B)** Representative images of GFP (All Foxp3) and RFP (Foxp3-FL only) single channels and overlay are shown (scale bar=150μm). **(C)** Representative image showing mRFP and eGFP co-expression (white arrow, Foxp3-FL Treg) or eGFP single expression (cyan arrow, Foxp3-ΔE2 Treg) (scale bar=5μm). **(D)** The CD3 (yellow) and MHC II (gray) single channels were used to identify and draw the T cell zone in each lymph node (scale bar=150μm). **(E)** Representative images of Foxp3-FL (red) and Foxp3-ΔE2 (cyan) Treg localization within and outside of the T cell zone (yellow dashed line) are shown for the non-draining LN (scale bar=150μm for whole LN and 20μm for zoom-in). *n*=3-5 mice. \**p*<0.05, \*\**p*<0.01, \*\*\**p*<0.001, \*\*\*\**p*<0.0001 by two-tailed paired t tests with Holm-Sidak method of correction for multiple comparisons as appropriate.

